# Evidence for an Organometallic Species Formed in the ArsL Reaction

**DOI:** 10.64898/2026.04.29.721666

**Authors:** Dante A. Serrano, Bo Wang, Squire J. Booker, Alexey Silakov

**Author notes:** **Corresponding author** Alexey Silakov, Squire J. Booker.

## Abstract

The radical *S*-adenosylmethionine (SAM) superfamily comprises over 800,000 unique sequences of enzymes that catalyze more than 100 distinct reactions. Canonical radical SAM (RS) enzymes are composed of a full or partial triose phosphate isomerase fold and contain a highly conserved CX_3_CX_2_C motif. The cysteines in the conserved motif ligate an [Fe_4_S_4_] cluster used in the reductive cleavage of SAM to yield methionine and a 5′-deoxyadenosyl 5′-radical (5′-dA•). The 5′-dA• is typically used to initiate catalysis by abstracting a hydrogen atom from a bound substrate, yielding 5′-deoxyadenosine and a substrate radical. ArsL, a recently characterized RS enzyme involved in the biosynthesis of the antibiotic arsinothricin (AST), catalyzes a reaction that deviates from the canonical RS reaction, forming methylthioadenosine and a 3-amino-3-carboxypropyl radical (ACP•). The ACP• is used to construct a carbon-arsenic bond to form the organoarsenic compound hydroxyarsinothricin (AST-OH), the penultimate enzymatic step in forming AST. While investigating how ArsL suppresses the formation of the 5′-dA• in favor of the ACP•, we discovered that ArsL forms a unique organometallic species containing a bond between the gamma carbon of ACP and the unique iron of the [Fe_4_S_4_] cluster. We propose that ArsL uses this novel species to generate a sufficiently electrophilic carbon that can be attacked by arsenous acid, thereby forming the carbon-arsenic bond.

## Introduction

Arsenic is a commonly occurring environmental toxin. This metalloid is often taken up by cells and mistaken for environmental phosphorus. To mitigate the adverse effects of arsenic, nature has evolved several pathways for arsenic detoxification, ranging from the direct methylation of arsenic by *S*-adenosylmethionine (SAM)-dependent methyltransferases for cellular export to more complex modifications [1]. The methylation reactions lead to the formation of monomethylarsenous acid (MMA(III)), dimethylarsenous acid (DMA(III)), and trimethylarsenous acid (TMA(III)), which all exhibit decreased arsenic reduction potentials, allowing it to be more readily oxidized by molecular oxygen to pentavalent As(V), which is less toxic than its trivalent As(III) counterpart [2-18].

An alternative pathway for arsenic detoxification involves incorporating arsenic into organic metabolites, some of which are bioactive. For example, bisenarsan and the speculative arsenic-containing fatty acids are secondary metabolites produced by Actinobacteria in response to environmental arsenic. These molecules are thought to form via an arsenoenolpyruvate intermediate, with the pyruvyl group from either the shikimate pathway or from pyruvate scavenged from the cellular environment being transferred to inorganic arsenate [19-28]. Radical *S*-adenosylmethionine (RS) enzymes also play a prominent role in generating arsenic-containing organic molecules. These enzymes, all of which contain at least one [Fe_4_S_4_] cluster, overwhelmingly catalyze the reductive cleavage of *S*-adenosylmethionine (SAM) to methionine and a 5′-deoxyadenosyl 5′-radical (5′-dA•). Most often, the 5′-dA• initiates catalysis by abstracting a hydrogen atom (H•) from the enzyme-bound substrate. One such RS enzyme, ArsS, plays a key role in the biosynthesis of various oxo-arsenosugars. It appends the 5′-dA• to an organo-arsenate molecule, generating 5′-deoxy-5′-dimethylarsinoyl-adenosine (DDMAA). The adenine of DDMAA can then be removed, allowing for further derivatization of the ribose [29-34].

ArsL, another RS enzyme, is involved in the biosynthesis of arsinothricin (AST), an arsenic-containing metabolite that displays antimicrobial activity [35-37]. AST resembles L-phosphinothricin, an herbicidal and antibacterial metabolite that inhibits glutamine synthetase.

While phosphinothricin biosynthesis requires 24 gene products, AST biosynthesis requires only two [38-40]. In the first step, catalyzed by ArsL, a 3-amino-3-carboxypropyl moiety from SAM is transferred to trivalent arsenic (As(III)) to afford hydroxyarsinothricin (AST-OH). This species is subsequently methylated by ArsM, then oxidized to As(V), forming AST [41-43].

Interestingly, in contrast to studies on ArsS, studies by several labs have shown that ArsL from *Burkholderia gladioli* reductively cleaves SAM to generate a 3-amino-3-carboxypropyl radical (ACP•) rather than the 5′-dA• generated by the overwhelming majority of RS enzymes [44-46]. This ACP• is believed to be an intermediate in the ArsL reaction; however, its exact mode of reactivity in generating AST-OH is unknown. In this work, we characterize an ArsL from *Pseudomonas aeruginosa*, which exhibits 60% sequence identity with that from *B. gladioli*. We show that the ACP• most likely forms an adduct with the unique iron of the [Fe_4_S_4_] cluster in ArsL to generate an [Fe_3_-S_4_-Fe-ACP]^3+^ species, reminiscent of the well-characterized species omega observed in a number of different RS enzymes [47-51]. We posit that this species is an intermediate on the pathway to forming the arsenic-carbon bond.

## Results

### Expression and characterization of *Pa*ArsL

The gene encoding *Pa*ArsL was cloned into a pET28a plasmid such that the encoded protein would contain an N-terminal hexahistidine (His_6_)-tag to enable its purification by immobilized metal affinity chromatography (IMAC). The construct was used to transform *Escherichia coli* BL21 (DE3) cells along with the pDB1282 and pBAD42-BtuCEDFB plasmids [52, 53]. The pDB1282 plasmid enhances iron-sulfur (FeS) cluster incorporation, while the pBAD42-BtuCEDFB plasmid, for reasons that we do not understand, enhances general protein solubility. The UV-vis spectrum of anaerobically purified *Pa*ArsL (**Figure S1**) shows the presence of a peak at ∼420 nm, with a spectral envelope and color (brown) that suggests the presence of an [Fe_4_S_4_] cluster. Consistent with this assignment, the purified protein contains 4.8 ± 0.7 Fe ions per monomer [54]. Moreover, when *Pa*ArsL is reduced with sodium dithionite (DT), it displays an electron paramagnetic resonance (EPR) spectrum consistent with an [Fe_4_S_4_]^1+^ cluster (**Figure S2B**).

### Elucidation of the *Pa*ArsL arsenic binding mode

A representative multiple-sequence alignment of predicted *Pa*ArsLs reveals six absolutely conserved cysteine residues (**Figure S3**). Three of these cysteines, C209, C213, and C216, ligate the [Fe_4_S_4_] cluster required for the reductive cleavage of SAM [44]. Three additional cysteines, C439, C440, and C443, lying in a CCXXC motif in the C-terminal region of the protein, are believed to be involved in arsenic binding [7-9, 55-57]. Studies by Dong and colleagues have shown that removal of this motif via the construction of a *Bg*ArsL_ΔC-Term_ variant results in loss of activity [45]. Further studies by Mauger et al. showed that individual substitutions of these cysteines with Ala also result in the loss of activity [44]. Herein, we use intact-protein mass spectrometry (IP-MS) to probe the role of these cysteines in arsenic binding. As-isolated (AI) *Pa*ArsL was denatured using a 10-fold excess of methanol, then pelleted and washed with water before resolubilizing it and analyzing it by IP-MS. AI wild-type (wt) *Pa*ArsL purifies with a fraction (30-54%) of adventitiously bound arsenic (**Figure 2A, red triangle**). Upon treating AI wt-*Pa*ArsL with 2.5 equivalents of arsenic, the protein becomes fully bound with at least one arsenic ion, as denoted by the 72.4 Da shift in the deconvoluted mass plot (**Figure S4**). Upon engineering a *Pa*ArsL triple variant containing Cys to Ala substitutions in the RCCLKC motif (*Pa*ArsL_AALKA_), we observe no arsenic signatures in the IP-MS spectrum of either the as-isolated protein or arsenic-treated protein, supporting the role of these cysteine residues in binding arsenic (**Figure 2B**). Whether arsenic is bound by all three cysteine residues or by only one or two is currently unknown.

**Figure 1:**
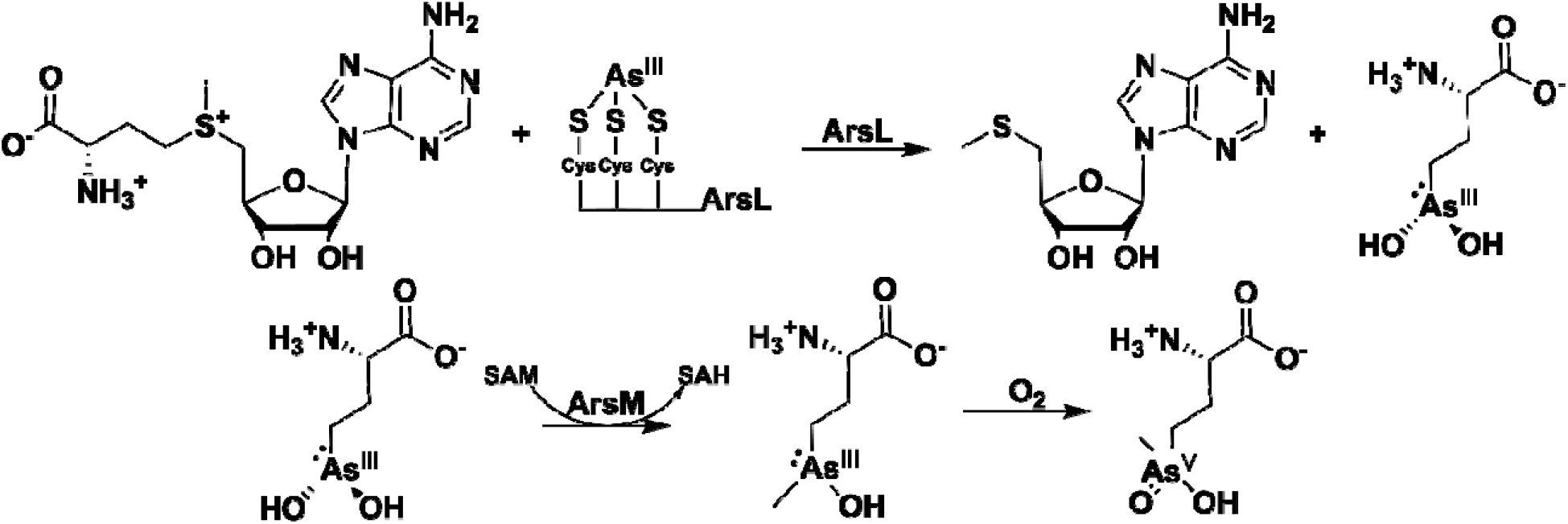
Proposed pathway for the biosynthesis of AST. The first enzyme in the reaction is ArsL, a canonical RS enzyme. ArsL appends a 3-amino-3-carboxypropyl moiety derived from SAM to arsenic to form hydroxyarsinothricin. This is then methylated by a canonical SAM-dependent arsenic methyltransferase, ArsM, forming trivalent arsinothricin. It is then proposed that the trivalent form undergoes non-enzymatic oxidation with molecular oxygen to form pentavalent arsinothricin, the final product.

**Figure 2:**
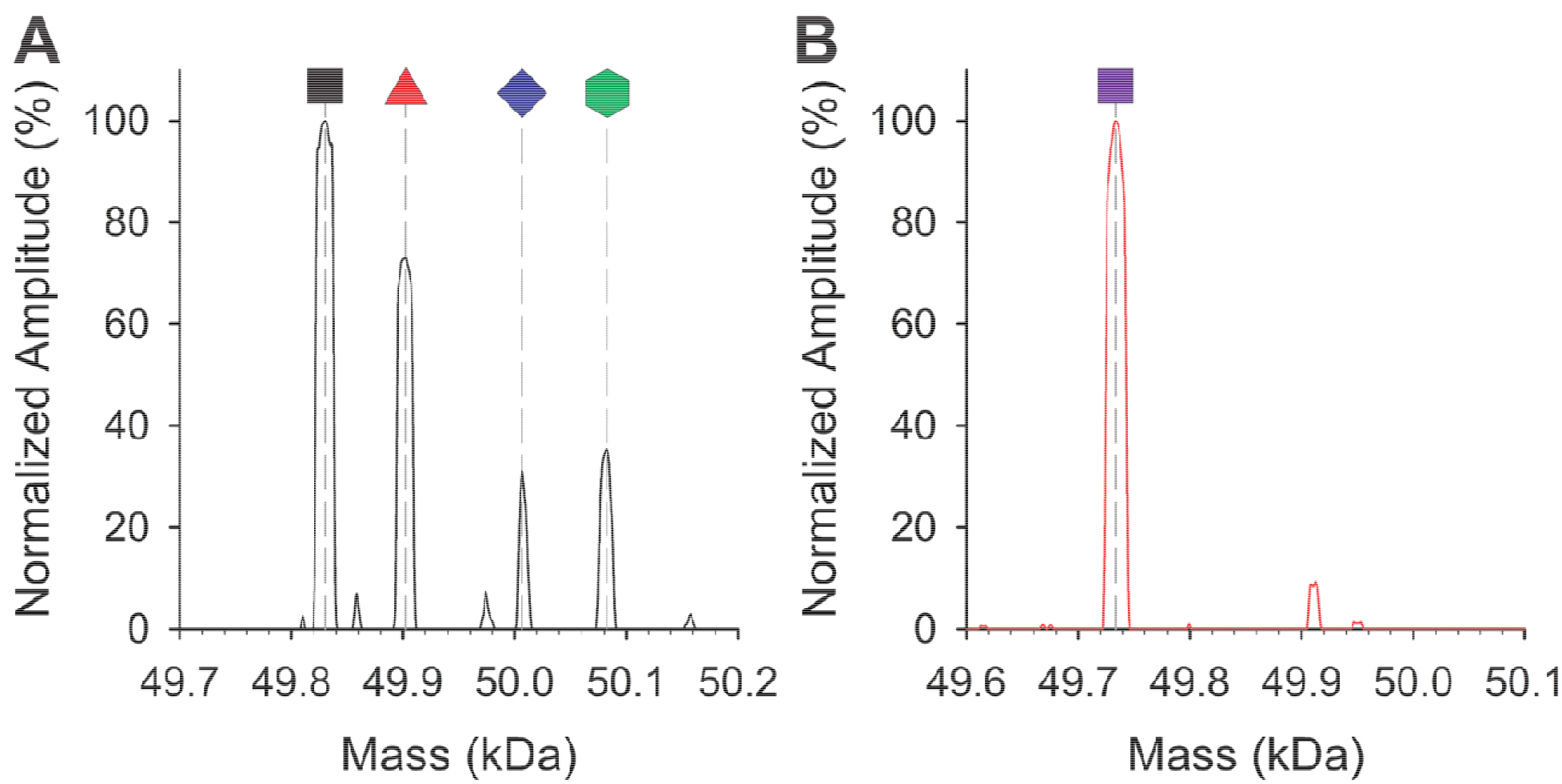
Arsenic binds to the C-terminal region of *Pa*ArsL. A) As-isolated *Pa*ArsL, when precipitated with methanol before being resolubilized in 1.5% Formic acid and 5% acetonitrile before being injected into the mass spectrometer, contains several species. Apo-*Pa*ArsL, 49,829.90 Da observed (black square), theoretical apo-*Pa*ArsL mass is 49,831.57 Da. *Pa*ArsL-As 49,902.30 Da observed (red triangle), theoretical *Pa*ArsL-arsenic mass is 49,903.47 Da. There are two other higher-mass species, 50,004.78 Da and 50,080.71 Da (blue diamond and green hexagon). B) The as-isolated *Pa*ArsL_AALKA_ variant precipitated and run with the same experimental conditions, there is only one species present, the apo-*Pa*ArsL_AALKA_, 49,733.45 Da observed (purple square), theoretical *Pa*ArsL_AALKA_ mass is 49,735.73 Da.

W*t-Pa*ArsL exhibits a 72.4 Da mass shift compared to the apo enzyme upon arsenic binding. The mass shift associated with adding arsenic to the fully protonated protein should be 74.922, suggesting that the shift also includes the loss of 2 (72.91 Da, -3.21 ppm) or 3 protons (71.89, - 23.44 ppm). We suggest that the observed mass shift corresponds to the loss of 3 protons, because arsenic would bear an additional OH group (89.91 Da, -343.95 ppm) if it were ligated by two cysteines in an aqueous environment rather than by three. High-resolution mass spectrometry (HRMS) was used to assess substrate binding to *Pa*ArsL. Samples were prepared with and without 2.5 equivalents of arsenic, pelleted, washed, and solubilized as before, and then fragmented using both data-directed and all-ion-fragmentation approaches. Upon fragmentation, we were able to collect peptide data at isotopic resolution (120,000), whereas the intact protein data were collected at 15,000. Fragmentation data from the control sample not treated with arsenic did not support the presence of bound arsenic using a confidence value of 21.53 ppm or ± 0.20 Da from the expected. Arsenic-treated wt-*Pa*ArsL showed evidence of arsenic binding by all three C-terminal cysteines. Arsenic-bound peptides displaying a 71.89 Da mass shift were resolved to 27.97 ppm or ± 0.34 Da from the expected. No fragments were detected that correlated with arsenic bound by two cysteines and no OH group. Moreover, fragments of arsenic bound by two cysteines with an OH group exceeded errors of 40 ppm, further supporting that arsenic is bound to *Pa*ArsL by all three terminal cysteines (**Figure S5**).

### Production of AST-OH

Several studies of *B. gladioli* ArsL have provided evidence for the products of the reaction with SAM and arsenic being MTA, ACP•, and AST-OH. Byproducts have also been observed, including 2-aminobutyric acid (2-ABA) and homocysteine sulfenic acid (HCSA). HCSA is formed when dithionite (DT) is used as the requisite low-potential reductant, presumably resulting from the recombination of the ACP• with the dithionite radical. By contrast, 2-ABA is formed when the ACP• obtains the equivalent of an H• ultimately from solvent [44, 45, 58]. In our study, we measured *Pa*ArsL’s activity under anoxic conditions at room temperature in reactions containing SAM (600 µM), sodium meta-arsenite (250 µM), and DT (750 µM). Reactions were removed from direct light before initiating with SAM and quenched at designated times. The quench solution contained hydrogen peroxide to oxidize the presumable AST(III)-OH species to the final AST(V)-OH species. Reaction mixtures were then analyzed by LC-MS using appropriate standards for quantification. **Figure 3A** shows the production of AST(V)-OH over a 2 h incubation, with 100 µM *Pa*ArsL catalyzing the formation of ∼23 µM AST-OH, indicating that the enzyme produces less than one turnover under the stated conditions. MTA forms in a burst, followed by a hyperbolic phase, although the SAM shows a small amount of MTA contamination. Similar to studies on *B. gladioli* ArsL, we also observe the formation of HCSA at concentrations exceeding (by less than a factor of 2) those of AST(V)-OH.

**Figure 3:**
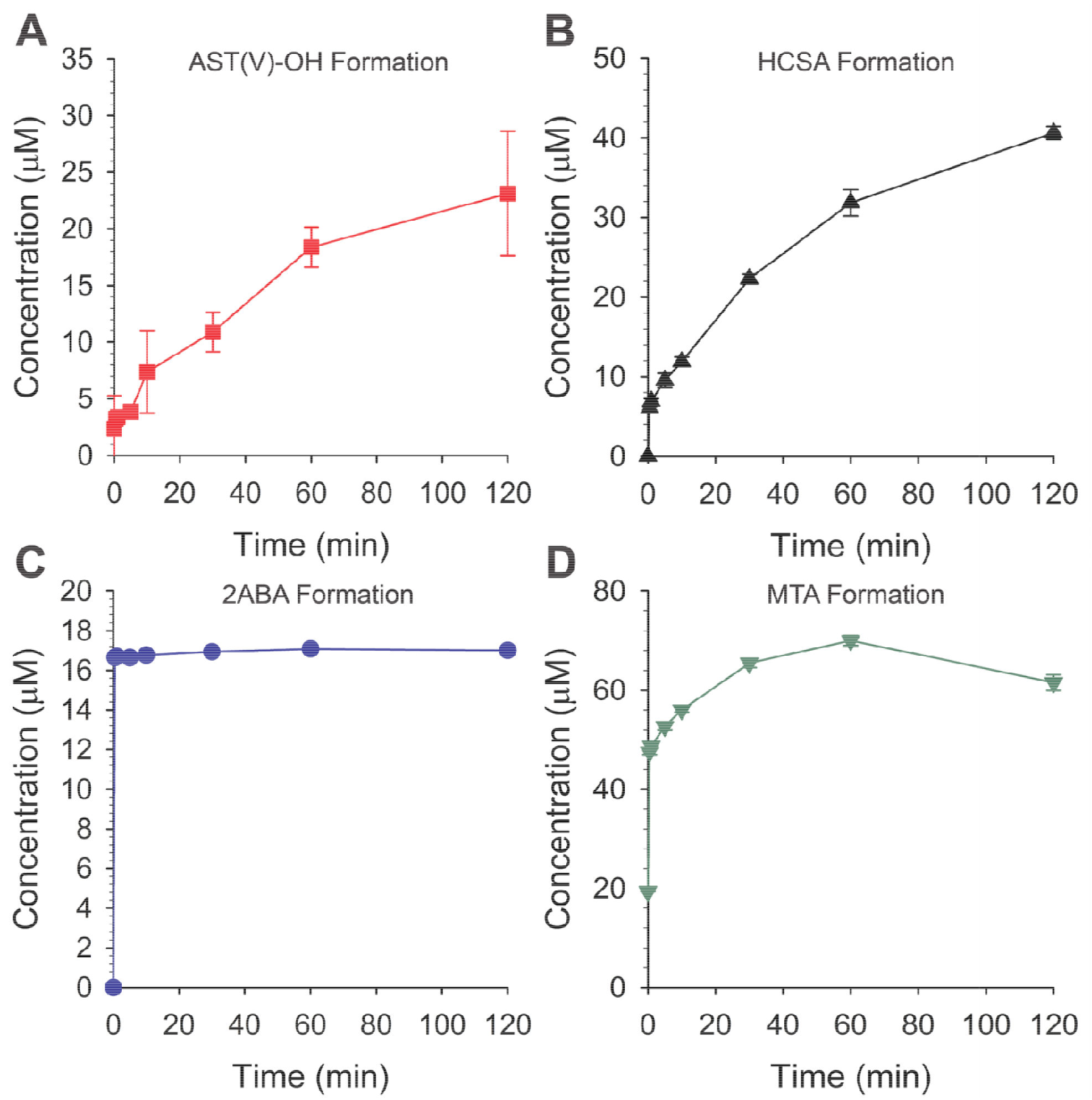
*Pa*ArsL catalyzes the formation of AST-OH from As(III) and SAM. A) *Pa*ArsL activity in the presence of DT, SAM, and As(III), quenched with H_2_O_2_ to a final concentration of 6%. Under these conditions, *Pa*ArsL generates AST(V)-OH over time. B) A portion of each reaction was not treated with H_2_O_2_, allowing quantification of ACP• and the [Fe_3_S_4_–Fe–ACP]^3+^ species intercepted by DT to form HSCA. C) The second nonproductive pathway monitored is solvent quenching of ACP• to form 2ABA, which appears only as an initial burst. D) MTA, the coproduct of SAM cleavage during ACP• formation, was also quantified. HC was monitored but remained below the limit of quantification. Error bars represent one standard deviation from triplicate reactions, with the mean shown at the center.

### Spectroscopic characterization of *Pa*ArsL

Initial characterization of *Pa*ArsL’s [Fe_4_S_4_] cluster was carried out using continuous-wave EPR (CW EPR) spectroscopy. As-isolated *Pa*ArsL displays a weak signal near g = 2.0 with the shape typical for an [Fe_3_S_4_]^1+^ cluster (**Figure S2A**). Upon reduction with DT, a strong axial spectrum, consistent with an [Fe_4_S_4_]^1+^ cluster, is observed with principal g-values of [2.029, 1.911, 1.893] (**Figure S2B**). When the sample is prepared with 300 µM *Pa*ArsL in the presence of 2 mM SAM, the EPR spectrum is completely lost within 10 min (**Figure S2D**). This loss is consistent with a reductive cleavage of SAM that results in the oxidation of the cluster to an EPR-silent [Fe_4_S_4_]^2+^ species, commonly observed for radical SAM enzymes [46]. Similarly prepared samples containing 300 µM *Pa*ArsL produce 146 ± 27 µM HCSA and 117 ± 12 µM MTA under the same conditions. Notably, when reduced *Pa*ArsL is incubated with *S*-adenosylhomocysteine (SAH), a SAM analog that cannot undergo reductive cleavage, the EPR signal persists. We note that the EPR signal bears a distinct rhombic distortion, presumably reflective of coordination of SAH to the [Fe_4_S_4_]^1+^ cluster [46] (**Figure S2C**).

### Characterization of SAM binding

To resolve whether a novel SAM-binding mode to the [Fe_4_S_4_] cluster is the cause of the non-canonical chemistry in *Pa*ArsL, we utilized hyperfine sublevel correlation (HYSCORE) spectroscopy to probe the chemical environment of the [Fe_4_S_4_]^1+^ cluster. To assign which of SAM’s functional groups are in close proximity to the [Fe_4_S_4_]^1+^ cluster, samples were prepared with SAM at natural abundance (na-SAM) or SAM containing various isotopic substitutions, including [*methyl*-^13^C]-SAM, [*carboxy*-^13^C]-SAM, [*amino*-^15^N]-SAM, and [^13^C_5_,-^15^N]-SAM. [^13^C_5_,-^15^N]-SAM is enriched with ^13^C at all methionine-derived carbons and ^15^N at the α-position. To prevent reductive cleavage of SAM upon binding to the [Fe_4_S_4_]^1+^ cluster, we prepared samples in the absence of reductant (i.e., in the [Fe_4_S_4_]^2+^-SAM form) and then subjected the samples to cryoreduction at 77 K with a 6 Mrad γ-irradiation dose, allowing detection of SAM binding to the [Fe_4_S_4_]^1+^ cluster [59]. We collected HYSCORE data at the maximum EPR absorption of the [Fe_4_S_4_]^1+^ cluster (g = 1.919), away from the radical signals around g = 2.002 that are formed as a byproduct of cryoreduction (**Figure S6**).

In the *Pa*ArsL sample with na-SAM, we observed a set of single- and double-quantum cross-correlation peaks characteristic of ^14^N (**Figure 4A**). The ^15^N-labeling of the amino group of SAM ([*amino*-^15^N]-SAM) shows a distinct modification of the HYSCORE spectra, unambiguously identifying the α-amino nitrogen of SAM as one of the coordinating ligands to the unique iron of the [Fe_4_S_4_] cluster (**Figure 4D**). HYSCORE spectra of *Pa*ArsL with bound [*carboxy*-^13^C]-SAM exhibit a noticeable cross-correlation ridge near the Larmor frequency of ^13^C (3.737 MHz at 349.0 mT). This weak coupling suggests the carboxy group of SAM is in close proximity to the [Fe_4_S_4_] cluster (**Figure 4C**). Additionally, samples prepared with [^13^C_5_,^15^N]-SAM show an increase in the general intensity of the weakly coupled ^13^C feature, as compared to ^14^N, ^15^N, and ^1^H signals, indicating an increase in the number of proximally located ^13^C atoms contributing to the signal. However, there is no additional feature or change in the breadth of the cross-correlation ridge, suggesting that the ^13^C-labeled carboxy carbon is likely the closest atom of SAM’s methionine group to the [Fe_4_S_4_] cluster. [^13^C_5_,^15^N]-SAM also produces the same ^15^N feature shift seen with [*amino*-^15^N] -SAM (**Figure 4E**). By contrast, samples prepared with [*methyl*-^13^C]-SAM show no significant deviation in HYSCORE spectra from that of the na-SAM sample (**Figure 4B**).

**Figure. 4:**
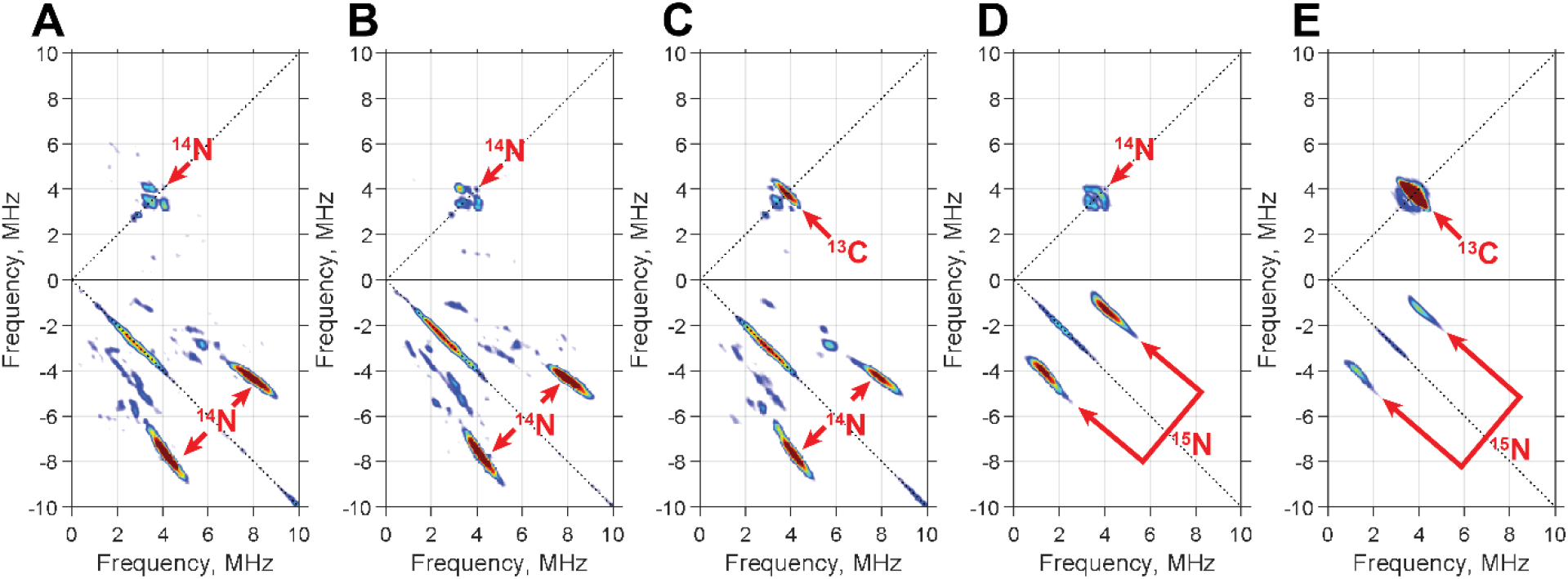
SAM binds to the reduced [Fe_4_S_4_]^1+^ cluster of *Pa*ArsL. *Pa*ArsL (500 µM) incubated with SAM (5 mM) or its isotopically labeled forms and cryoreduced before HYSCORE analysis. A) Natural abundance SAM serves as the control. B) [*methyl*-^13^C]-SAM shows no perturbation in the spectra when compared to the control. C) [*carboxy*-^13^C]-SAM shows a weak hyperfine correlation at the Larmor frequency of ^13^C. D) [*amino*-^15^N]-SAM shows a strong hyperfine correlation at twice the Larmor frequency of ^15^N. E) [^13^C_5_, ^15^N]-SAM shows the same spectral shifts as both [*carboxy*-^13^C]-SAM and [*amino*-^15^N]-SAM, but with a higher intensity of the ^13^C cross-correlation ridge. Samples were measured at 10 K; MW frequency 9.5 - 9.37 GHz, τ = 140 ns, and [π/2] = 8 ns.

When the experimental data for SAM binding to *Pa*ArsL are compared with previously collected HYSCORE measurements for the RS enzyme RlmN and previously-reported hyperfine coupling constants for SAM-bound [Fe_4_S_4_] clusters (**Figure S8C and S8D**), our results show little to no deviation [60, 61]. Consequently, this data unequivocally shows that *Pa*ArsL positions SAM similarly to the prototypical SAM-binding mode of canonical RS enzymes, which normally favors selective cleavage of the γ-carbon over the 5′-carbon. This outcome is surprising given that formation of 5′-deoxyadenosine was not observed in our assays, nor was it reported for *B. gladioli* ArsL [44, 45, 62]. We also note that Dph2, which also catalyzes an unconventional SAM cleavage, has a distinct SAM-binding mode compared to canonical RS enzymes (**Figure S7**)[58].

### Characterization of an [Fe_4_S_4_]^3+^-like cluster

When *Pa*ArsL samples are prepared under turnover conditions, we observe an EPR spectrum distinct from that of the [Fe_4_S_4_]^1+^ cluster, whether in the presence or absence of SAM or SAH, or from that of the as-isolated protein. The EPR spectrum observed during turnover is best fitted to one major and two minor species. The major species (∼80% of signal intensity) exhibits g-values of 2.074, 2.004, and 1.989, while one unknown minor species (∼18% of signal intensity) exhibits g-values of 2.130, 2.023, and 2.005. The remaining minor species, representing an [Fe_4_S_4_]^1+^ cluster, exhibits g-values of 2.059, 1.930, and 1.925 (**Figure 5**). The major signal is markedly different from those typically observed for [Fe_4_S_4_]^1+^ clusters. The g >2 principal components of this species are more in line with those for [Fe_4_S_4_]^3+^ and organometallic species observed in other FeS cluster-containing enzymes (**Table S2**). The identity of the minor species remains elusive and will require further investigations. In this study, we focus on the dominant spectral component. To assess the nature of major species, we enriched *Pa*ArsL with ^57^Fe by overproducing it in *E. coli* containing ^57^Fe in the growth media and incubated it with an excess of [^13^C_5_,^15^N]-SAM. To avoid an adverse reaction with sodium dithionite and the formation of HCSA, the protein was reduced using a stoichiometric concentration of titanium(III) citrate. The EPR signal of the ^57^Fe-enriched sample exhibits a distinct broadening compared to the sample containing natural-abundance iron, suggesting that the signal emanates from the FeS cluster. The inclusion of four ^57^Fe nuclei with a hyperfine coupling constant averaging ∼33 MHz allowed for an accurate reproduction of the ^57^Fe-induced line broadening, confirming that this species originates from an [Fe_4_S_4_] cluster. The EPR simulation shown in **Figure 5** was aided by ^57^Fe electron-nuclear double resonance (ENDOR) spectroscopy (**Figure S9**), which was used to estimate the ^57^Fe hyperfine coupling constants. These measurements, in conjunction with the characteristic principal g-values, indicate that the major signal is due to an [Fe_4_S_4_]^3+^ cofactor.

**Figure 5:**
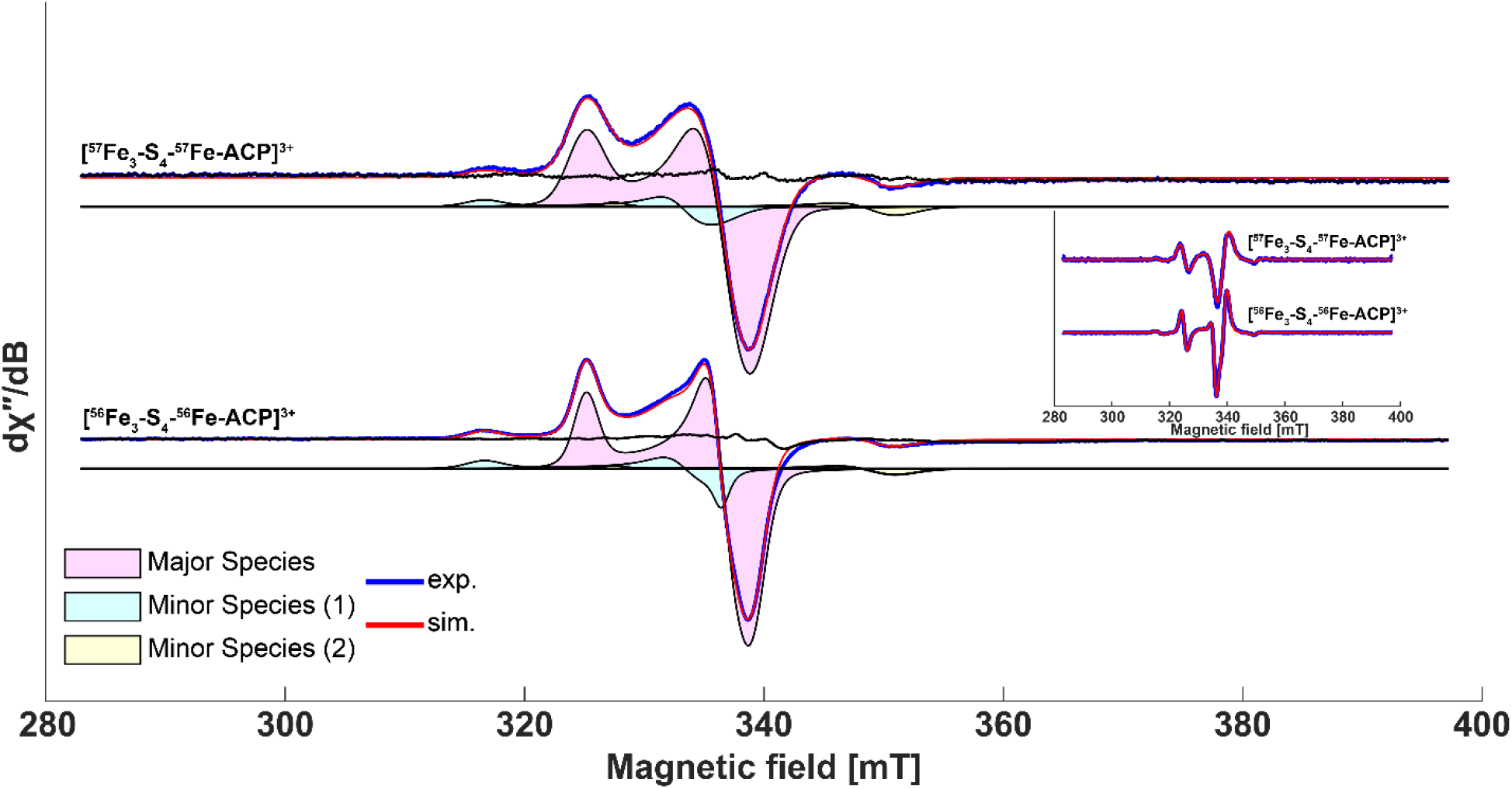
CW X-Band EPR of the ^56^Fe and ^57^Fe labeled organometallic species. The [Fe_3_-S_4_-Fe-ACP]^3+^ spectra were collected using CW EPR (blue) and simulated with three species. The major species has g-values of [2.074, 2.004, and 1.989], and the two minor species have g-values of [2.130, 2.023, 2.005], and [2.059, 1.930, 1.925]. All individual components are shown as shaded areas, with the sum of their intensities shown in red and overlaid on the experimental data. The difference between the experimental data (blue) and the simulation (red) is shown as a solid black line. The [^57^Fe_3_-S_4_-^57^Fe-ACP]^3+^ spectra are shown in blue. The simulation of the major species from the ^56^Fe data included the hyperfine couplings of four ^57^Fe nuclei, with the minor species unchanged; the sum of their intensities is shown in red, and the difference from the experimental data is shown in black. The ^57^Fe hyperfine coupling from each nucleus to best simulate the broadening seen in the data are Fe_1_ & _2_ = [29, 29, 29] MHz and Fe_3_ & _4_ = [42.35, 42.35, 28.93] MHz. The inset shows the pseudomodulated spectrum of experimental and simulated data, further demonstrating that the simulation agrees with the experimental data.

The observation that the [Fe_4_S_4_]^3+^ cluster is generated under the reducing conditions required for the reductive cleavage of SAM suggests that it might derive from an organometallic species containing an Fe-carbon bond. We therefore examined the samples by field-dependent HYSCORE spectroscopy at Q-band frequencies. As shown in **Figure 6**, these measurements reveal an exceptionally strong ^13^C hyperfine coupling when the sample is prepared with [^13^C_5_,^15^N]-SAM. Although the hyperfine coupling shows a slight field dependence across the magnetic field range, the observed 18-23 MHz interaction is fully consistent with an Fe–C bond, supporting assignment of the species as an [Fe_3_-S_4_-Fe-ACP]^3+^ cluster [47-51, 61, 63-71]. These HYSCORE spectra also reveal distinct ^15^N cross-correlation signals, suggesting that the amino group of the ACP remains coordinated to the cluster after SAM cleavage.

**Figure 6:**
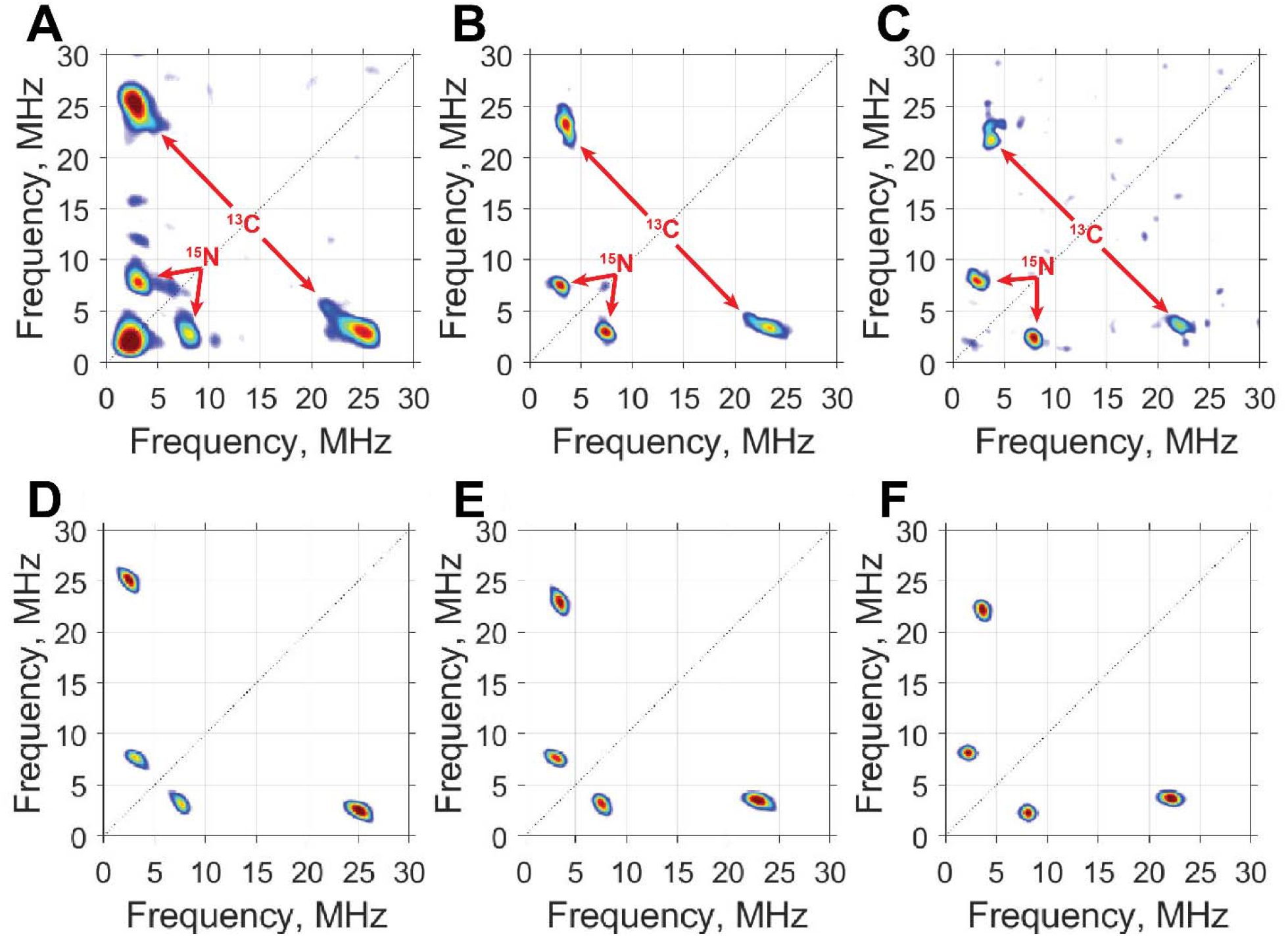
Q-band HYSCORE of [Fe_3_-S_4_-Fe-ACP]^3+^ prepared with [^13^C_5_,^15^N]-SAM demonstrating ACP coordination by a Fe-^13^Cγ bond and the coordination of the ^15^N-labeled amino group. A-C) Experimental data measured at (A) 1211 mT, (B) 1194 mT, and (C) 1176.5 mT. D-F) Are simulations consistent with experimental data in A-C. ^13^C hyperfine coupling is simulated to be [27.5, 18.2, 17.0] MHz with an orientation of [0°, 9°, 30°] (y-convention Euler angles). The weakly coupled ^15^N was simulated to be [6.5, 5.8, 0.5] MHz with an orientation of [0°, 90°, 0°] (y-convention Euler angles). Samples were measured at 10 K; MW frequency 34.006 GHz, τ = 172 ns, and [π/2] = 12 ns.

Additionally, we collected Q-band HYSCORE spectra with ^57^Fe-enriched *Pa*ArsL in the presence of [*carboxy*-^13^C]-SAM. The absence of an observable ^13^C cross-correlation ridge in the HYSCORE spectrum for this sample indicates that the carboxylate group of the ACP moiety is not coordinated to the unique iron of the [Fe_3_-S_4_-Fe-ACP]^3+^ cluster (**Figure S10**). Overall, HYSCORE spectra demonstrate that the ACP is coordinated to the unique site of the FeS cluster by its γ-carbon and amino group.

**Scheme. 1:**
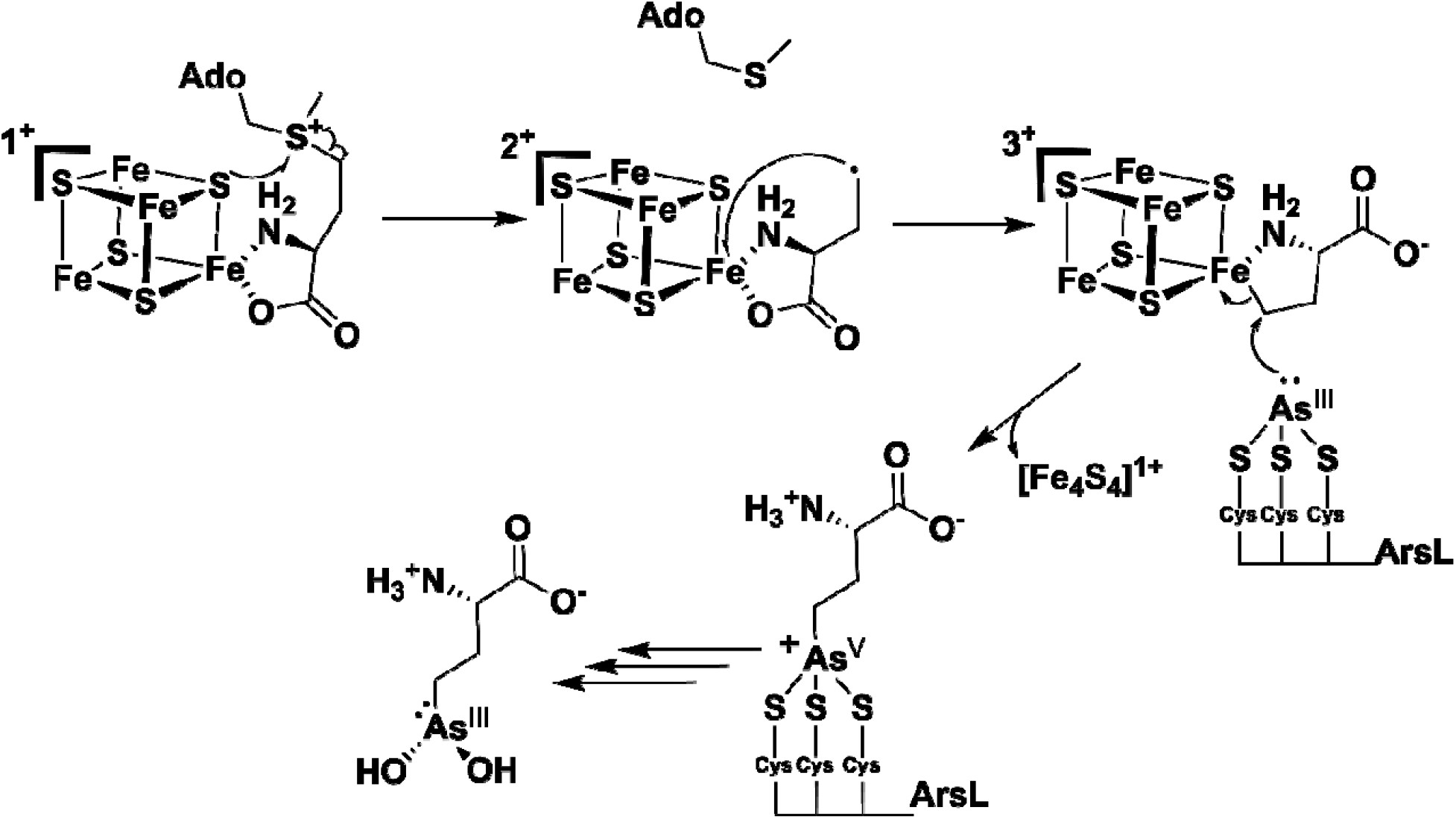
Proposed ArsL reaction mechanism. After the reductive cleavage of SAM to form the ACP•, the radical adds to an Fe of the [Fe_4_S_4_]^2+^ cluster to form a [Fe_3_-S_4_-Fe-ACP]^3+^ species, activating the γ-carbon of the ACP moiety for the subsequent nucleophilic attack from bound arsenous acid to form the product-bound AST-OH. We suggest that the arsenic is coordinated by three conserved cysteine residues on the protein.

## DISCUSSION

RS enzymes, particularly those within the Radical SAM superfamily (Pfam: PF04055), overwhelmingly catalyze the reductive cleavage of SAM to generate methionine and a 5′-dA•. The uniqueness of ArsL is that it is a canonical RS enzyme but does not show typical RS reactivity. Outside of ArsL, the only other known outlier is tryptophan methyltransferase, TsrM, which does not reductively cleave SAM at all and instead performs cobalamin-based polar chemistry [60, 72, 73]. One additional enzyme, Dph2, involved in diphthamide biosynthesis, catalyzes the reductive cleavage of SAM to generate an ACP• [74]. However, Dph2 is not a canonical RS enzyme. It lacks the characteristic partial TIM-barrel fold and therefore does not reside within PF04055. Moreover, it binds SAM in a conformation that is distinct from that of canonical RS enzymes (**Figure S6**). These characteristics have been used to rationalize its noncanonical reductive cleavage of SAM. Conversely, *Pa*ArsL falls squarely within PF04055 both in sequence and in structure, and our HYSCORE analysis of the cryoreduced enzyme shows that SAM binds its [Fe_4_S_4_] cluster in a bidentate fashion through its α-amino and α-carboxylate moieties like other characterized RS enzymes in PF04055.

Under turnover conditions, but in the absence of arsenic, *Pa*ArsL accumulates a distinctly different EPR-active species. The spectrum of this species is centered at a g_avg_ > 2.0, suggesting an [Fe_4_S_4_]^3+^-like cluster. Broadening of the EPR spectrum of the ^57^Fe-enriched *Pa*ArsL confirms that it arises from an FeS cluster. Moreover, the inclusion of four ^57^Fe nuclei was necessary to simulate the line-broadening in the EPR spectrum appropriately, confirming that the species is an [Fe_4_S_4_]^3+^-like cluster. Importantly, formation of this species requires both SAM and a low-potential reductant. This suggests that the [Fe_4_S_4_]^3+^-like cluster is generated when the ACP• from SAM’s reductive cleavage adds to an iron ion of the [Fe_4_S_4_]^2+^ cluster, leading to a 1e^-^ oxidation of the cluster and resulting in the [Fe_4_S_4_]^3+^-like signal. The formation of organometallic species has been previously observed in RS enzymes, with the species termed Ω considered an intermediate in the formation of the 5′-dA• in canonical RS enzymes. An Ω-like species, termed Intermediate I, has also been observed in Dph2 [50, 58]. Moreover, organometallic species have been proposed in non-RS enzyme reactions such as IspG and IspH; however, their structures have not been unambiguously established. To determine the structure of the organometallic species in *Pa*ArsL, we prepared samples of the [Fe_4_S_4_]^3+^-like cluster with [^13^C_5_, ^15^N]-SAM and analyzed them by field-dependent HYSCORE spectroscopy. Analysis and simulation of the spectra revealed that ACP’s amino nitrogen is weakly coupled to the cluster. We also identified a single, unusually strong ^13^C_γ_ coupling to the cluster, with no evidence of a second, weaker coupling. The large magnitude of this carbon hyperfine interaction (21 MHz) is consistent with an Fe-C bond, while the absence of a weaker coupling suggests that ACP’s carboxyl group is no longer coordinated to the iron.

We do not believe that *Pa*ArsL’s organometallic species is similar to Ω in canonical RS enzymes or Intermediate I in Dph2. *Pa*ArsL binds SAM like traditional RS enzymes but cleaves the S-C_γ_ bond of SAM rather than the S-C5′ bond. The structure of Intermediate I has not been elucidated in detail, but can be represented by any of the five possibilities shown in **Figure 7**. As detailed above, our spectroscopic studies support species 2 as the structure for *Pa*ArsL’s [Fe_3_-S_4_-Fe-ACP]^3+^ organometallic species. Hyperfine coupling constants for Fe-C bonds in Ω of pyruvate formate-lyase activating enzyme and Intermediate I in Dph2 have been reported at 9.4 and 7.8 MHz, respectively. By contrast, the hyperfine coupling constant of the Fe-C bond in the [Fe_3_-S_4_-Fe-ACP]^3+^ organometallic species of *Pa*ArsL is significantly larger (21 MHz). Why the coupling constant in *Pa*ArsL differs so dramatically from those in Ω and Intermediate I remains unclear. One possibility is that they may reflect how each of these species reacts. In Ω and Intermediate I, the Fe-C bond undergoes homolysis to allow the carbon to react as a radical. By contrast, in *Pa*ArsL, we propose that the organometallic species may react electrophilically, allowing cleavage of the Fe-C bond by a polar mechanism upon nucleophilic attack by the arsenic atom.

**Figure 7:**
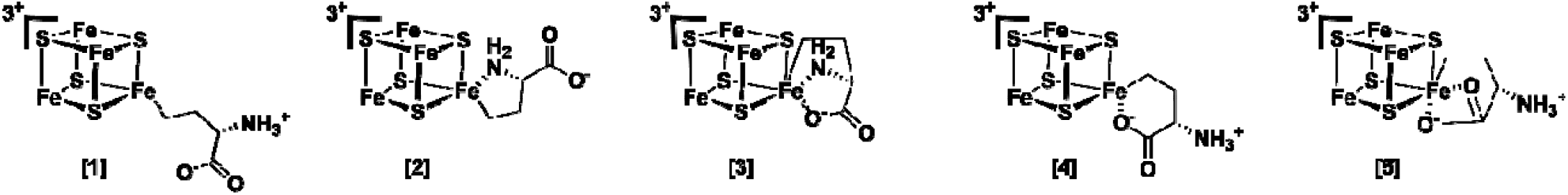
Proposed structures for the [Fe_3_-S_4_-Fe-ACP]^3+^ species.

At present, we favor the mechanism in Scheme 1 as the most consistent with our experimental observations. However, due to limitations of delivering arsenic to the active site of *Pa*ArsL after generating the organometallic species, we were unable to observe the direct conversion of the organometallic species to AST-OH. Therefore, we also offer an alternative. The Fe-C bond could undergo homolysis, with the nascent carbon radical attacking the bound arsenic. The formation of the carbon-arsenic bond would require the loss of an electron from the arsenic, which might be used to reduce the resulting [Fe_4_S_4_]^2+^ cluster to the [Fe_4_S_4_]^+^ state. Further studies are necessary to distinguish between these two possibilities.

## Materials and Methods

### Materials

Kanamycin, ampicillin, spectinomycin, and isopropyl β-*D*-1-thiogalactopyranoside (IPTG) were purchased from Gold Biotechnology (St. Louis, MO). *N*-(2-Hydroxyethyl)piperazine-*N*′-2-ethanesulfonic acid (HEPES), tris(2-carboxyethyl)phosphine (TCEP), 2-mercaptoethanol (BME), phenylmethylsulfonyl fluoride (PMSF), SIGMA FAST EDTA-Free protease inhibitor tablet, sodium *meta*-arsenite, and sodium dithionite were from Sigma Corp (St. Louis, MO). Ferric chloride was from EMD Biosciences (Gibbstown, NJ), while Coomassie brilliant blue was from ICN Biomedicals (Aurora, OH). The Bradford reagent for protein concentration determination and the bovine serum albumin standard were purchased from Pierce, Thermofisher Scientific (Rockford, IL). ^57^Fe metal (98%) was obtained from Cambridge Isotope Laboratories (Tewksbury, MA).

### Synthesis of SAM, isotopically labeled SAM molecules, Ti(III) citrate and AST-OH

*S,S*-SAM was synthesized from ATP and L-methionine using SAM synthetase and purified as described previously [75]. *S*-adenosyl[^13^C-*methyl*]methionine was synthesized similarly using [^13^C-*methyl*]methionine, while *S*-adenosyl[^15^N-*amino*]methionine, *S*-adenosyl[^13^C-*carboxy*]methionine, and *S*-adenosyl[^13^C_5_,^15^N-*amino*]methionine were synthesized similarly using [^15^N-*amino*]methionine, [^13^C-*carboxy*]methionine, [^13^C_5_,^15^N-*amino*]methionine. Ti(III) citrate was prepared as described previously [76]. AST-OH was synthesized and purified as described previously [77].

### Protein mass spectrometry

Samples were prepared by precipitating 8.1 µmols of protein with a 1:10 volumetric ratio of methanol. Samples were centrifuged at 15,000 × g, and the supernatant was discarded. The pellet was resolubilized with 50 µL of a solution of 1.5% formic acid and 5% acetonitrile; samples were once again centrifuged at 15,000 x g to remove any unsolubilized protein. The solute was applied onto a BioBasic™ 4 HPLC column connected to a Thermo QE-HFX mass spectrometer for analysis at a resolution of 15,000. The data was then processed with UniDec. Fragments of As-bound *Pa*ArsL were generated with both data-directed fragmentation and allion-fragmentation approaches and collected at a resolution of 120,000. Fragments were deconvoluted with Thermo Freestyle and processed in ProSight Lite [78, 79].

### Expression and Purification of *Pa*ArsL

The *PaarsL* gene was cloned into a pET28a vector, which was used, along with plasmids pDB1282 [53] and pBAD42-BtuCEDFB [52], to transform *E. coli* BL21 DE3. The construct produces *Pa*ArsL containing a N-terminal hexahistidine (His_6_) tag. For expression, *E. coli* containing the appropriate plasmids was cultured at 37°C with shaking (165 rpm) in M9-ethanolamine minimal media as described previously [52]. At an OD_600_ of 0.3, L-cysteine (50 µM) and FeCl_3_ (100 µM) or ^57^FeCl_3_ (100 µM) were added to the culture, along with L-(+)-arabinose, required for induction of genes on the pDB1282 and pBAD42-BtuCEDFB plasmids. At an OD_600_ of 0.5, IPTG (200 µM) was added to induce the expression of *Pa*ArsL, which was allowed to continue at 18°C for 16 h. Cells were then harvested by centrifugation at 6,000 × g for 15 min at 4 °C. Purification of *Pa*ArsL, variants, and ^57^Fe-labeled *Pa*ArsL was carried out in a Coy (GrassLake, MI) anaerobic chamber (≤1 ppm O_2_). Frozen cell paste was thawed in lysis buffer (50 mM HEPES, pH 7.5, 450 mM KCl, 4 mM imidazole, 10 mM β-mercaptoethanol, 10 mM MgCl_2_, 10% (v/v) glycerol, 1 mg/mL lysozyme, 0.5 mg/mL DNase, 15 µg/mL PMSF, and a SIGMA FAST EDTA-Free protease inhibitor tablet). The cells were lysed by sonication (Fisher Scientific Model 550 sonic dismembrator) at 30% amplitude (6 pulses each for 45 s with 5 min between each pulse) and then clarified by centrifugation at 55,000 × g for 1 h at 4 °C., The clarified lysate was loaded onto Ni-NTA resin, washed with 100 mL of wash buffer (50 mM HEPES pH 7.5, 450 mM KCl, 10 mM imidazole, 10 mM β-mercaptoethanol, 10 mM MgCl_2_, and 10% (v/v) glycerol), then eluted with elution buffer (50 mM HEPES, pH 7.5, 45 0mM KCl, 500 mM imidazole, 10 mM β-mercaptoethanol, 10 mM MgCl_2_, and 10% (v/v) glycerol). During elution, about 40 mL of dark brown eluant was collected, concentrated, and exchanged into gel filtration buffer (50 mM HEPES, pH 7.5, 200 mM KCl, 5 mM TCEP, 10%(v/v) glycerol) using a PE-10 column (Cytiva). The as-isolated *Pa*ArsL was concentrated again and immediately separated from aggregates by size-exclusion chromatography (Sephadex S200 26/60 HR column) using an AKTA (Cytiva) purification system housed in the anaerobic chamber. Yields were determined with Bradford analysis, using a correction factor of 0.645, as previously described [54].

### *Pa*ArsL activity assays

*Pa*ArsL activity assays were performed at room temperature (25.2 C°) under anaerobic conditions in a final volume of 1.2 mL. Each reaction contained 100 µM *Pa*ArsL, 50 mM HEPES (pH 7.5), 200 µM L-tryptophan (internal MS standard), 750 µM sodium dithionite, 200 mM KCl, and 10% (v/v) glycerol. After incubating for 10 min, 250 µM sodium meta-arsenite is added. After incubating further for 20 min, reactions are initiated by adding SAM to a final concentration of 600 µM.

Triplicate reactions were prepared and quenched using two separate quenching protocols. Each assay was split into two equal portions at the time of quenching. For analysis of MTA, HCSA, and 2ABA, samples were quenched with an equal volume of a solution containing 1% formic acid, 99% methanol, and 5 mM TCEP. For analysis of AST-OH, samples were quenched with an equal volume of a solution containing 1% formic acid in 99% methanol and subsequently treated with hydrogen peroxide to a final concentration of 6% (v/v).

Prior to LC–MS analysis, all samples were centrifuged at 14,000 × g for 10 min at 4 °C to remove precipitated protein, and the clarified supernatant was transferred to LC-MS vials. Chromatographic separation was performed on a ZORBAX RRHD SB-CN column (2.1 × 150 mm, 1.8 µm) equilibrated in 98% solvent A (0.1% formic acid in water) and 2% solvent B (0.1% formic acid in acetonitrile). Solvent B was held at 2% for 2 min, then increased linearly to 12.5% over 4 min, and then increased again to 60% over an additional 4 min before being increased to 98% over 2 min. The column was held at 98% B for 2 min, returned to 2% B over 1 min, and equilibrated at 2% B for an additional 2 min. Mass analysis was performed on an Agilent Technologies 1290 Infinity II series UHPLC system coupled to a 6470 QQQ Agilent Jet Stream electrospray-ionization mass spectrometer, and data were processed with Agilent MassHunter Quantitative Analysis 10.1 software [80].

### EPR sample preparation for cryoreduction experiments

Under anaerobic conditions, *Pa*ArsL (500 µM) in 50 mM HEPES (pH 7.5), 200 mM KCl, 5 mM TCEP, and 10% (v/v) glycerol was incubated for 10 min with 5 mM SAM, SAM isotopologs ([*amino*-^15^N]-SAM, [*carboxy*-^13^C]-SAM, or ^1^ [^13^C_5_, ^15^N]-SAM). Following incubation, samples were transferred into EPR tubes with 3.8mm O.D. and 2.78mm I.D. and flash-frozen in liquid nitrogen. Frozen samples were cryoreduced using the γ-irradiation facility at the Breazeale Nuclear Reactor (The Pennsylvania State University). EPR tubes were maintained in liquid nitrogen during irradiation and exposed to a total dose of 7.1 MRad from a ^60^Co γ-ray source, corresponding to an effective absorbed dose of approximately 6 MRad per sample [81].

### EPR sample preparation of the [Fe_4_S_4_]^3+^-like cluster

Under anaerobic conditions, *Pa*ArsL (850 µM) or ^57^Fe-labeled *Pa*ArsL (884 µM) in) in 50 mM HEPES (pH 7.5), 200 mM KCl, 5 mM TCEP, and 10% (v/v) glycerol was incubated with one reducing equivalent of titanium(III) citrate for 5 min. Reactions were then treated with six equivalents of SAM or SAM isotopologs ([*carboxy*-^13^C]-SAM, or [^13^C_5_, ^15^N]-SAM) and incubated for 55 min in the absence of light. Following incubation, samples were transferred via capillary under low-intensity red light into Q-band EPR tubes (2.8mm O.D. and 1.8mm I.D.) and flash-frozen in liquid nitrogen.

### EPR sample preparation of the [Fe_4_S_4_]^1+^ cluster

Under anaerobic conditions, *Pa*ArsL (300 µM) in 50 mM HEPES (pH 7.5), 200 mM KCl, 5 mM TCEP, and 10% (v/v) glycerol was incubated with 1mM of sodium dithionite for 10 min. Reactions were then treated with 2 mM of SAM or SAH. Following incubation, samples were transferred into EPR tubes with 4 mm O.D. and 3 mm I.D. and flash-frozen in liquid nitrogen. Data processing and simulation were performed in MATLAB 2020b using Kazan Viewer 2.0 and functions from EasySpin 5.2.36 [82-84].

### Continuous-wave electron paramagnetic resonance (CW-EPR)

EPR spectra were collected on a Magnettech MS5000X spectrometer equipped with an Oxford Instruments ESR900 continuous-flow liquid helium cryostat and a LakeShore 335 temperature controller. EPR spectra were recorded at a 50 µW microwave power from a range of 280 mT to 400 mT with a sweep time of 72 s, 100 kHz modulation frequency, and a 1 mT modulation amplitude. Data processing and simulation were performed in MATLAB 2020b using Kazan Viewer 2.0 and functions from EasySpin 5.2.36 [82-84].

### Pulse EPR measurements

All pulse EPR measurements were conducted on a Bruker Elexsys E580 spectrometer equipped with a SuperX-FT microwave bridge and an in-house constructed intermediate-frequency extension fitted with a Millitech 5W pulse power amplifier for Q-band measurements [85]. Cryogenic temperatures were reached using an Oxford CF935 helium-flow cryostat controlled by an Oxford Instruments ITC502 temperature controller. X-band measurements were carried out using a Bruker ER 4118X-MS5 resonator with a 1 kW traveling-wave tube (TWT) microwave amplifier (Applied Systems Engineering, model 117x). Q-band pulse EPR measurements were conducted on a TE011 resonator built in-house, utilizing the open resonator concept [85]. Q-band ENDOR measurements were performed on the same instrument. Radiofrequency was controlled by a Bruker DICE RF synthesizer and amplified by a BT01000-AlphaSA TOMCO 1kW amplifier. Data acquisition and control of experimental parameters were performed using Bruker XEPR software. Processing of the data and simulations of the HYSCORE and ENDOR data were performed in MATLAB 2020b using Kazan Viewer 2.0 and functions from EasySpin 5.2.36[82-84]. The following pulse sequences were used: Free Induction decay: [π/2]_MW_-t_deadtime_-(detection), 2-pulse ESEEM: [π/2]_MW_-τ-[π]_MW_-τ-(detection), HYSCORE: [π/2]_MW_-τ-[π/2]_MW_-T1-[π]_MW_-T2-[π/2]_MW_-(detection), and Davies ENDOR: [π]_MW_-[π]_RF_-[π/2]_MW_-τ-[π]_MW_-(detection).

## Acknowledgement

We thank Zachary Van Horn from the Breazeale Nuclear Reactor of the Pennsylvania State University for assistance with the cryoreduction experiments. This work is partially based upon work supported by the National Science Foundation (CHE-1943748 to A.S.), National Science Foundation Graduate Research Fellowship Program (DGE-1255832), US National Institute of Health (GM122595 to S.J.B), and the Eberly Family Distinguished Chair in Science (to S.J.B.).

S.J.B. is an investigator of the Howard Hughes Medical Institute (HHMI). Any opinions, findings, and conclusions or recommendations expressed in this material are those of the author(s) and do not necessarily reflect the views of any of the agencies that have funded this work.

## Associated content

### Supporting Information

Supporting figures (S1-S10), supporting tables S1 and S2, and scheme S1. The following files are available free of charge.

## Author information

## Author Contributions

The manuscript was written through the contributions of all authors. All authors have given approval to the final version of the manuscript.

