## Supplemental Information for "Evidence for an Organometallic Species Formed in the ArsL Reaction"

**Table of contents**

**Page 2.** Table of contents

**Page 3.** Figure S1. UV-vis spectrum of as-isolated PaArsL

**Page 4.** Figure S2. CW-EPR of PaArsL cluster

**Page 5.** Figure S3. Two conserved cysteine-containing regions in AST-OH synthases

**Page 6.** Figure S4. Apo-PaArsL converts to arsenic-bound PaArsL when treated with arsenic

**Page 7**. Figure S5. Fragments of arsenic-bound PaArsL confirm the identity and location of arsenic

**Page 8.** Table S1. Intact protein species

**Page 9.** Figure S6. X-Band CW spectra of cryoreduced PaArsL with SAM

**Page 10.** Figure S7. Comparison of SAM binding in canonical and non-canonical RS enzymes

**Page 11.** Figure S8. SAM binding simulations

**Page 12.** Figure S9. ^57^Fe Davies ENDOR of the [Fe_3_-S_4_-Fe-ACP]^3+^ species

**Page 13.** Figure S10. HYSCORE of [Fe_3_-S_4_-Fe-ACP]^3+^ with [carboxy-^13^C]-SAM

**Page 14.** Table S2. FeS Cofactor reference table

**Page 15.** Scheme S1. Alternative ArsL reaction mechanism

**Page 16.** References

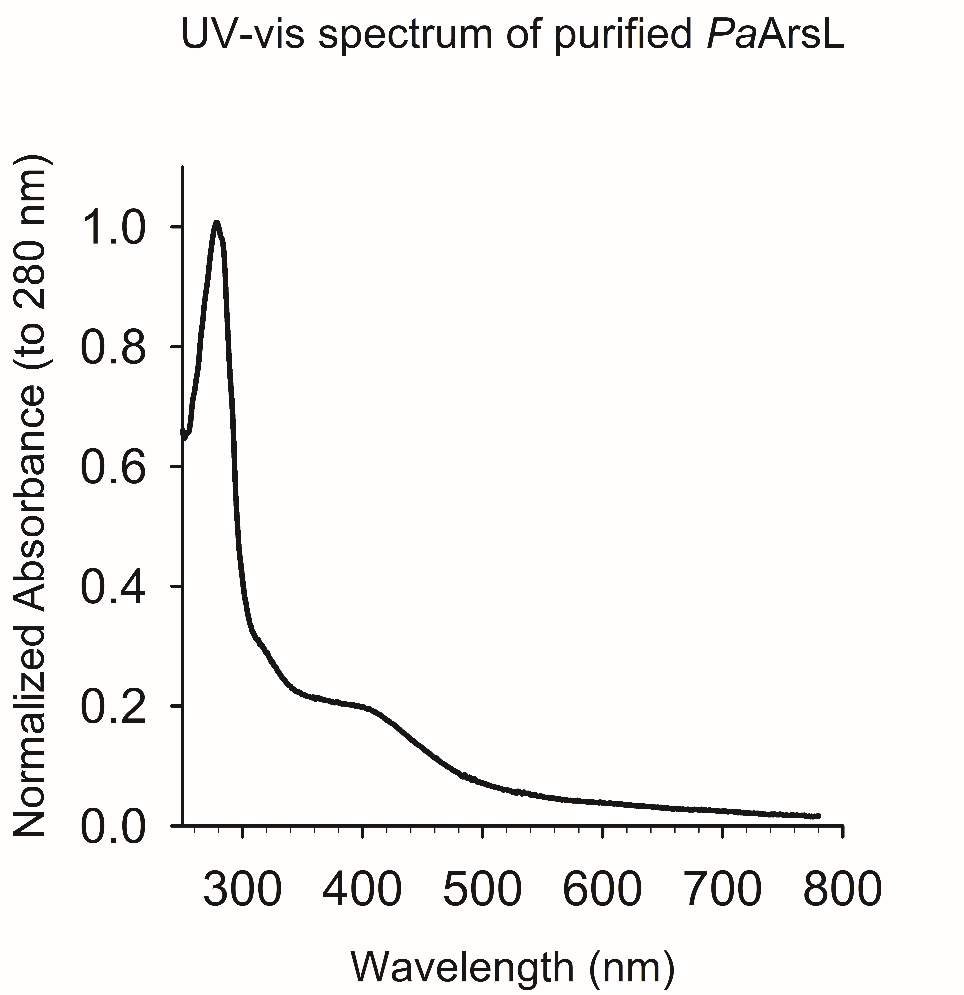

**Figure. S1: As-isolated *Pa*ArsL.** The UV-vis spectrum of anaerobically purified *Pa*ArsL has a prominent peak at ~420 nm, which plateaus before being dominated by the strong 280 nm feature of the protein.

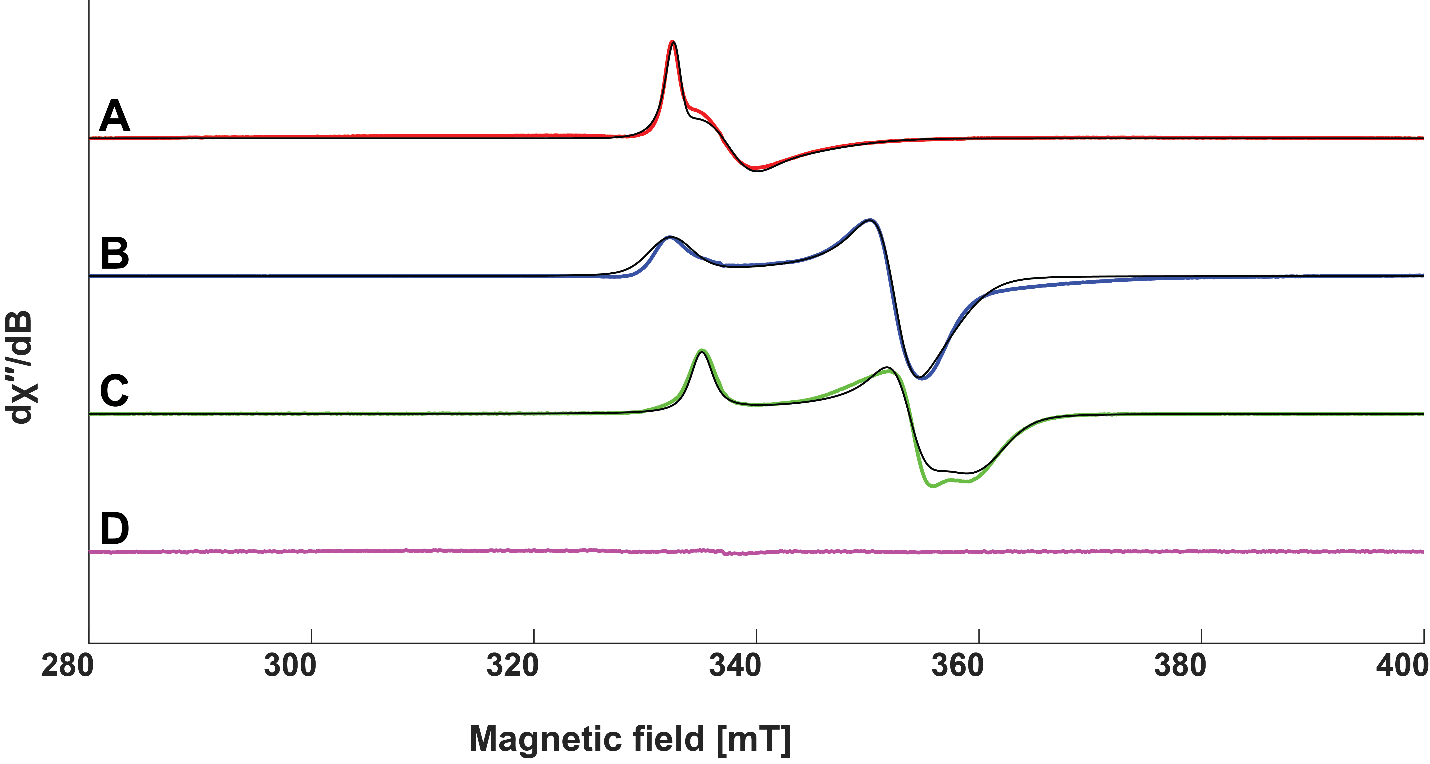

**Figure S2: CW-EPR of *Pa*ArsL cluster.** A) Red line is the experimental data of As-isolated *Pa*ArsL, which shows *a* [Fe_3_S^4^]^1+^ cluster *that was simulated in black and* yield g-values of [2.025, 1.993, 1.960], *Pa*ArsL (300 µM) in 50 mM HEPES (pH 7.5), 200 mM KCl, 5 mM TCEP, and 10% (v/v) glycerol. B) Blue line is the experimental data, the reduced [Fe_4_S_4_]^1+^ cluster of *Pa*ArsL. The simulation of this cluster (black) produced g-values of [2.029, 1.911, 1.893], *Pa*ArsL (300 µM) in 50 mM HEPES (pH 7.5), 200 mM KCl, 5 mM TCEP, and 10% (v/v) glycerol with 1 mM sodium dithionite. C) Green line is the experimental data *S*-adenosyl-homocysteine bound to the [Fe_4_S_4_]^1+^ cluster of *Pa*ArsL. A simulation of this cluster (black) produced g-values of [2.011, 1.905, 1.872], *Pa*ArsL (300 µM) in 50 mM HEPES (pH 7.5), 200 mM KCl, 5 mM TCEP, and 10% (v/v) glycerol with 2mM SAH and 1 mM sodium dithionite. D) The magenta trace is the EPR spectrum of *Pa*ArsL after being incubated with SAM for 8 seconds. *Pa*ArsL (300 µM) in 50 mM HEPES (pH 7.5), 200 mM KCl, 5 mM TCEP, and 10% (v/v) glycerol with 2 mM SAM and 1 mM sodium dithionite. All EPR spectra were recorded at a 50 µW microwave power from a range of 280 mT to 400 mT with a sweep time of 72 s, 100 kHz modulation frequency, and a 1 mT modulation amplitude.

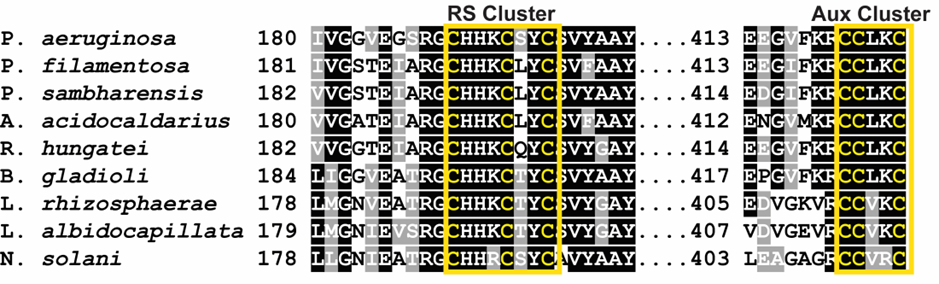

**Figure. S3: There are two conserved cystine regions in AST-OH synthases.** The first CX_3_CX_2_C motif is used to bind an [Fe_4_S_4_] cluster. The second CCX_2_C motif is used to bind the substrate, arsenous acid.

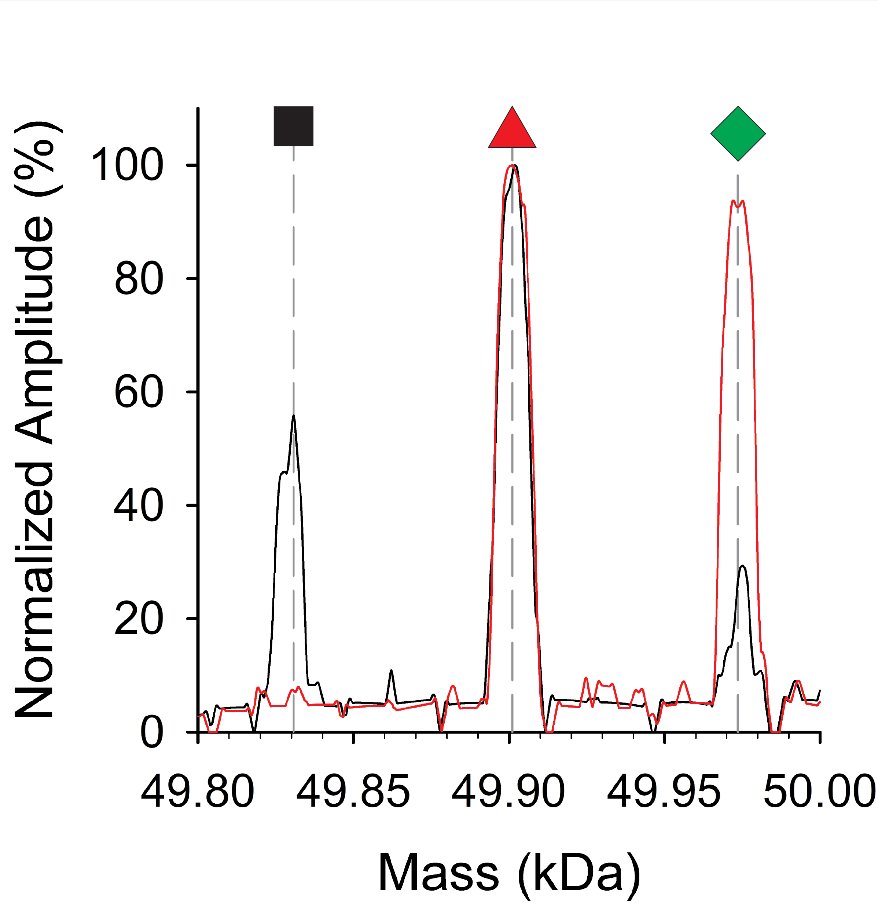

**Figure. S4: Apo-PaArsL converts to PaArsL-As when treated with arsenic.** Black line) As-isolated PaArsL before being treated with three equivalents of arsenic, has an observed mass of 49,830.67 Da observed (black square) theoretical apo-PaArsL mass is 49,831.57 Da. Red line) After being treated with arsenic all apo-PaArsL is converted into PaArsL-As with an observed mass of 49,901.00 Da observed (red triangle) and a theoretical mass of 49,903.47 Da. There is an additional species, caused by the overloading of arsenic onto PaArsL, observed at 49,973.65 Da. This species, PaArsL-As_2_, best correlates with PaArsL-As, with a second arsenic advantageously bound; it would have a theoretical mass of 49,975.37 Da.

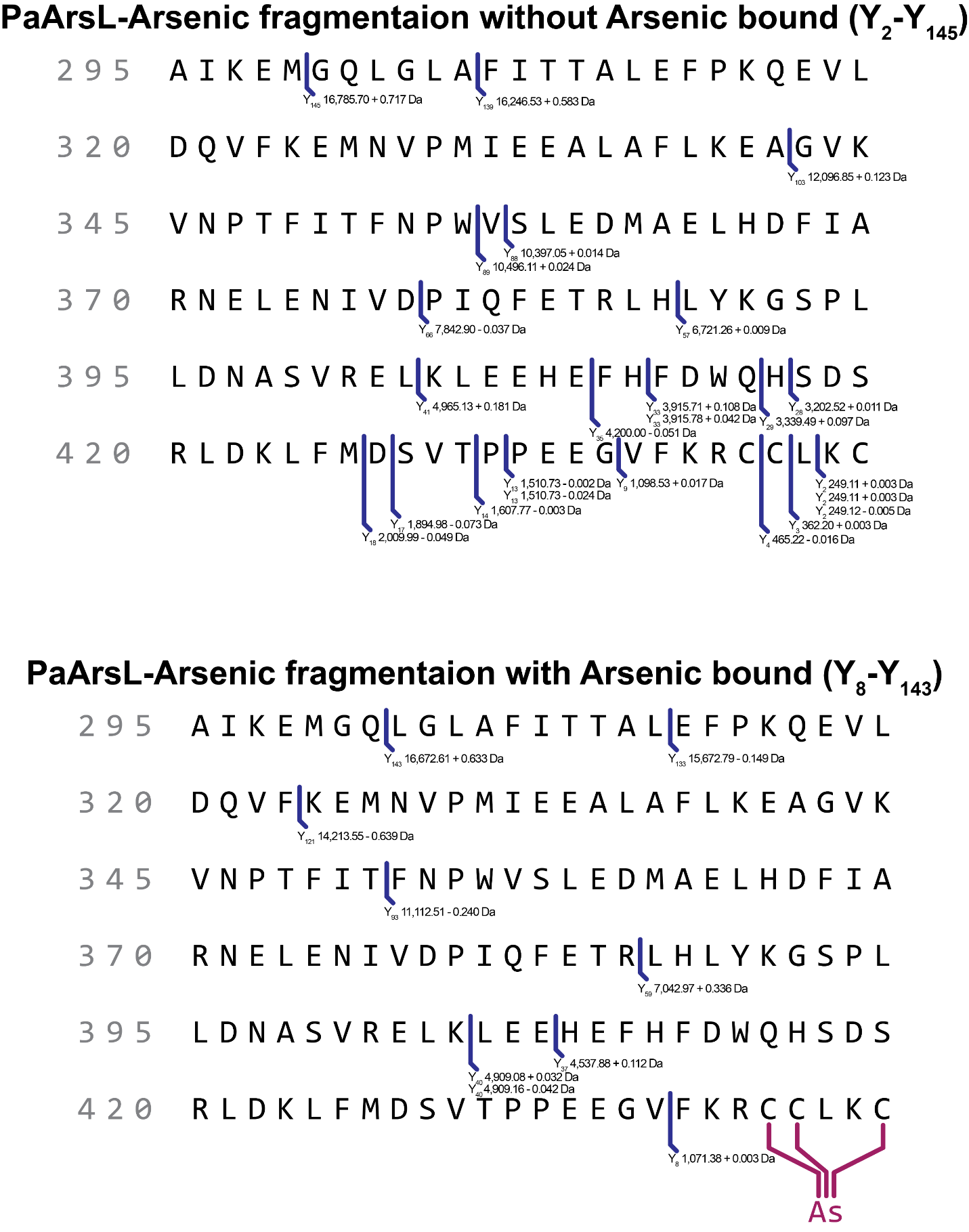

**Figure. S5: Fragments of the PaArsL-As confirm the identity and location of the modification**. PaArsL-As was fragmented by data-directed and all ion fragmentation methods. The fragments were then extracted from the run and deconvoluted to neutral masses and mapped to PaArsL with the use of ProSite Lite. Fragments without the arsenic added to the C-terminus provided a confidence of 21.53 RMS (ppm) or ± 0.20 RMS (Da), while Fragments with the arsenic added to the C-terminus provided a confidence of 27.97RMS (ppm) or ± 0.34 RMS (Da).

| **Symbol** | **Species** | **Structure** | **Observed mass (Da)** | **Theoretical mass (Da)** |
| --- | --- | --- | --- | --- |
| 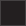 | Apo-*Pa*ArsL | 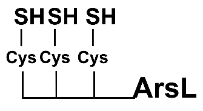 | 49,829.90 Da | 49,831.57 Da |
| 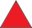 | *Pa*ArsL-As | 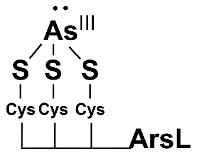 | 49,902.30 Da | 49,903.47 Da |
| 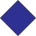 | *Pa*ArsL-AST-OH | 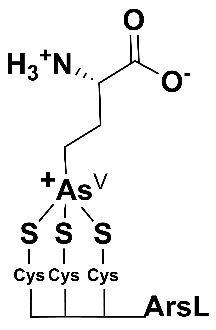 | 50,004.78 Da | 50,004.51 |
| 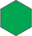 | *Pa*ArsL-AST-OH +advantageous arsenic | 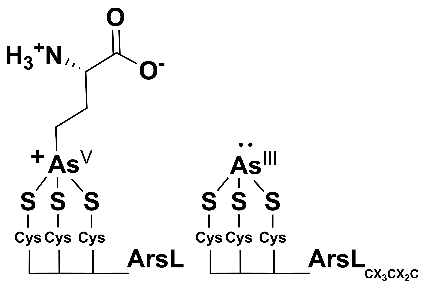 | 50,080.71 Da | 50,076.68 Da |
| 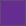 | apo-*Pa*ArsL_AALKA_ | 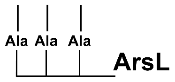 | 49,733.45 Da | 49,735.73 Da |

**Table. S1: Intact protein species**

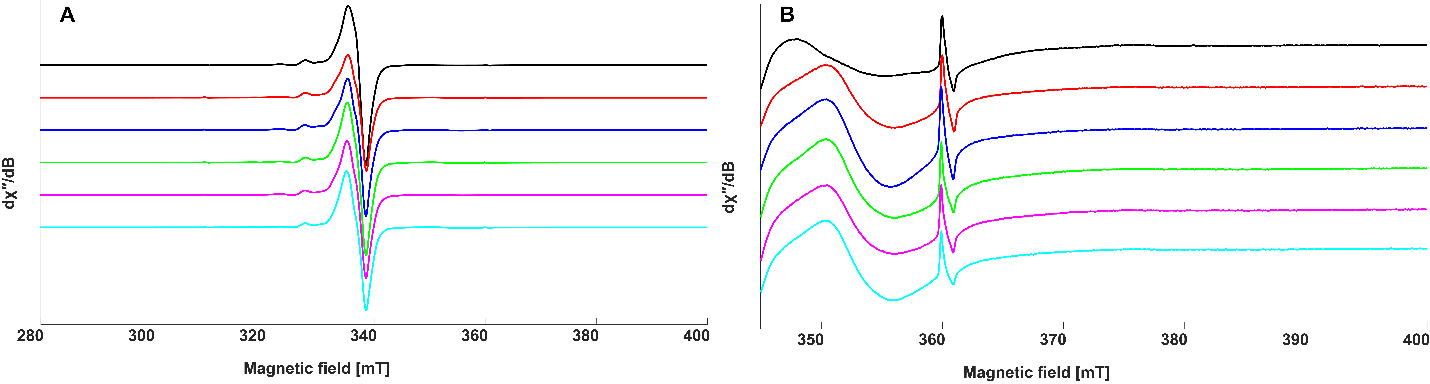

**Figure S6: X-Band CW spectra of the cryoreduced *Pa*ArsL with SAM samples.** A) Full spectra of the of cryoreduced *Pa*ArsL (500 µM) in 50 mM HEPES (pH 7.5), 200 mM KCl, 5 mM TCEP, and 10% (v/v) glycerol with 5 mM SAM or SAM isotopologs. B) Zoomed-in spectra of the cryoreduced *Pa*ArsL SAM samples. Black line is *Pa*ArsL without SAM, red line is *Pa*ArsL with SAM, blue line is *Pa*ArsL with [*methyl*-^13^C]-SAM, green line is *Pa*ArsL with [*carboxy*-^13^C]-SAM, magenta line is *Pa*ArsL with [^13^C_5_, ^15^N]-SAM, and cyan line is *Pa*ArsL with [*amino*-^15^N]-SAM. EPR spectra were recorded at a 50 µW microwave power from a range of 280 mT to 400 mT with a sweep time of 72 s, 100 kHz modulation frequency, and a 1 mT modulation amplitude.

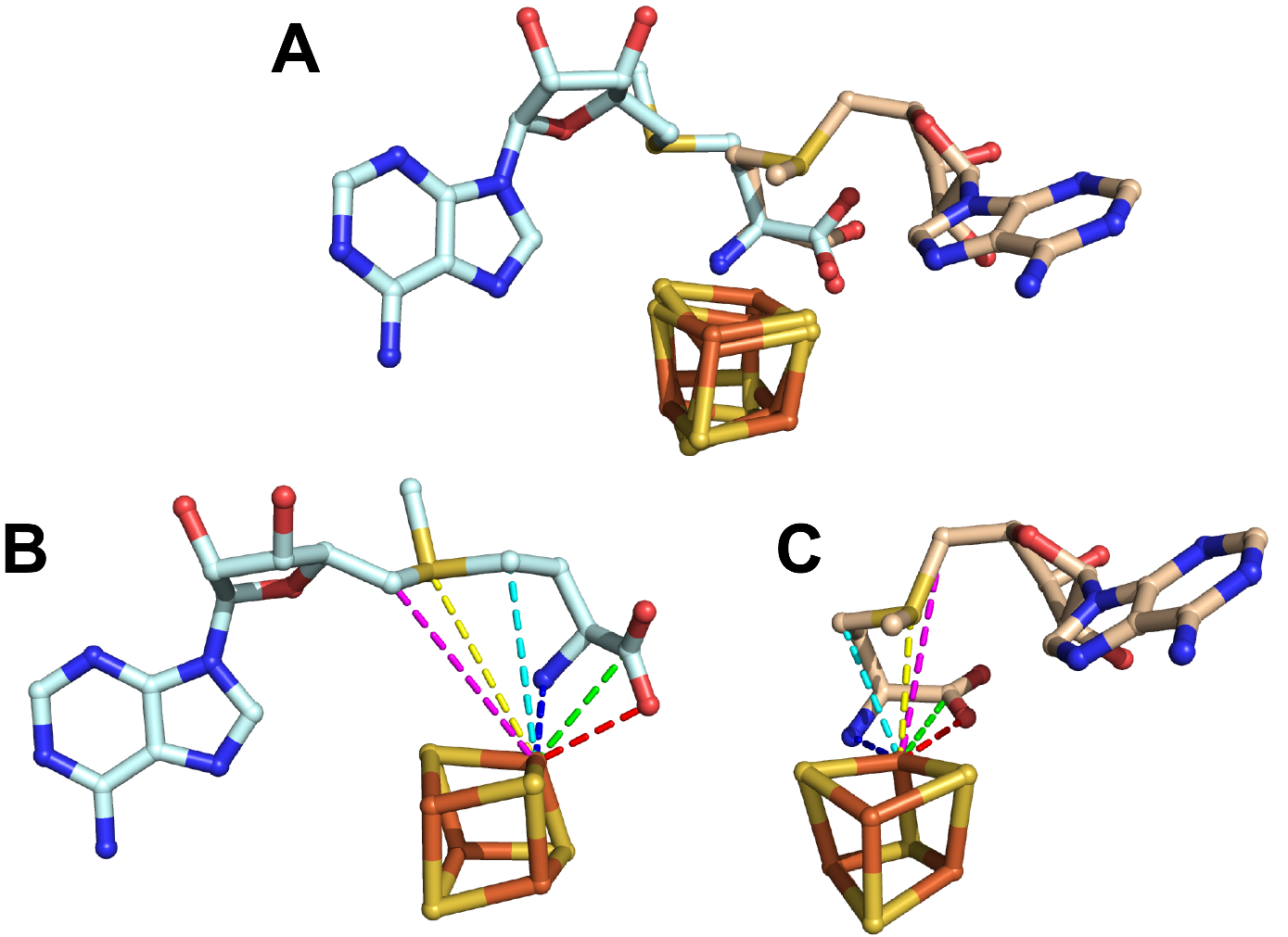

**Figure. S7: Comparison of SAM binding in canonical and non-canonical RS enzymes.** A) Overlay of SAM binding to the [Fe_4_S_4_] in *Ec*PFL-AE (PDB 3CB8)in tan and *Cmn*Dph2 (PDB 6BXN) in pale-blue. This shows that the non-canonical RS enzymes *Cmn*Dph2 bind SAM in a unique conformation when compared to *Ec*PFL-AE, a canonical RS enzyme. This is further confirmed by the differences in the differences of various atoms of SAM in relation to the cluster. B) The dashed lines signify the following distances of each atom to the unique iron of the cluster in *Cmn*Dph2: (red) carboxy-oxygen 2.7 Å, (green) carboxy-carbon 3.1 Å, (blue) amino-nitrogen 2.3 Å, (cyan) Cγ of SAM 4.3 Å, (yellow) sulfur of the sulfonium 4.7 Å, and (magenta) 5′-C of SAM 4.9 Å. C) The same measurements for *Ec*PFL-AE are: 2.1, 2.9, 2.2, 3.4, 3.2, and 5.0 Å.

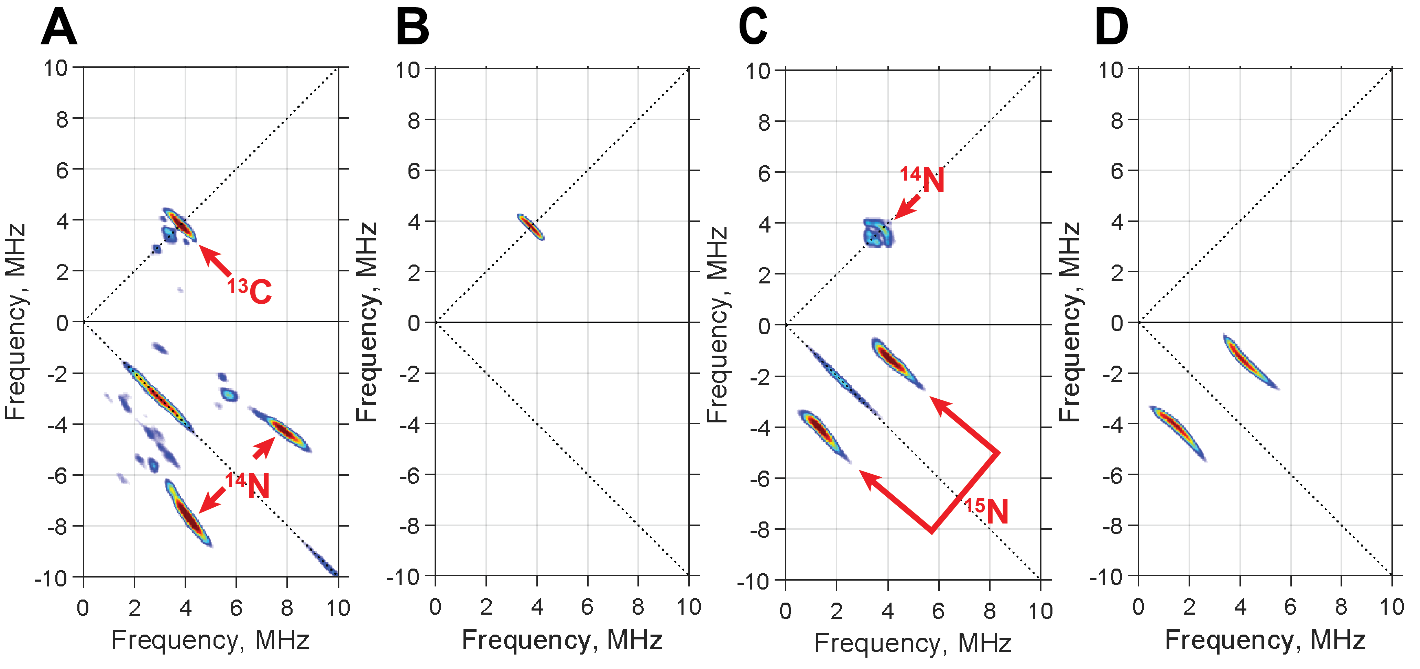

**Figure. S8: SAM binding simulations.** *Pa*ArsL (500 µM) incubated with SAM (5 mM) or its isotopically labeled forms and cryoreduced before HYSCORE analysis. A) [*carboxy*-^13^C]-SAM experimental data. B) Comparison of experimental data of [*carboxy*-^13^C]-SAM binding to the [Fe_4_S_4_] cluster of *Pa*ArsL with simulations using hyperfine parameters of canonical SAM binding is nearly indistinguishable The hyperfine interaction is simulated to yield an A tensor of [1.1, -0.95, -0.82] MHz and an orientation of [60°, 0°, 45°] (y-convention Euler angles). C) [*amino*-^15^N]-SAM experimental data. Samples were measured at 10 K; MW frequency 9.5 - 9.37 GHz, τ = 140 ns, and [π/2] = 8 ns. D) A similar conclusion is reached for [*amino*-^15^N]-SAM binding to the cluster. The hyperfine simulation of the amino group of SAM yields an A tensor of [9.7, 6, 3.5] MHz and an orientation of [75°, 75°, 90°] (y-convention Euler angles).

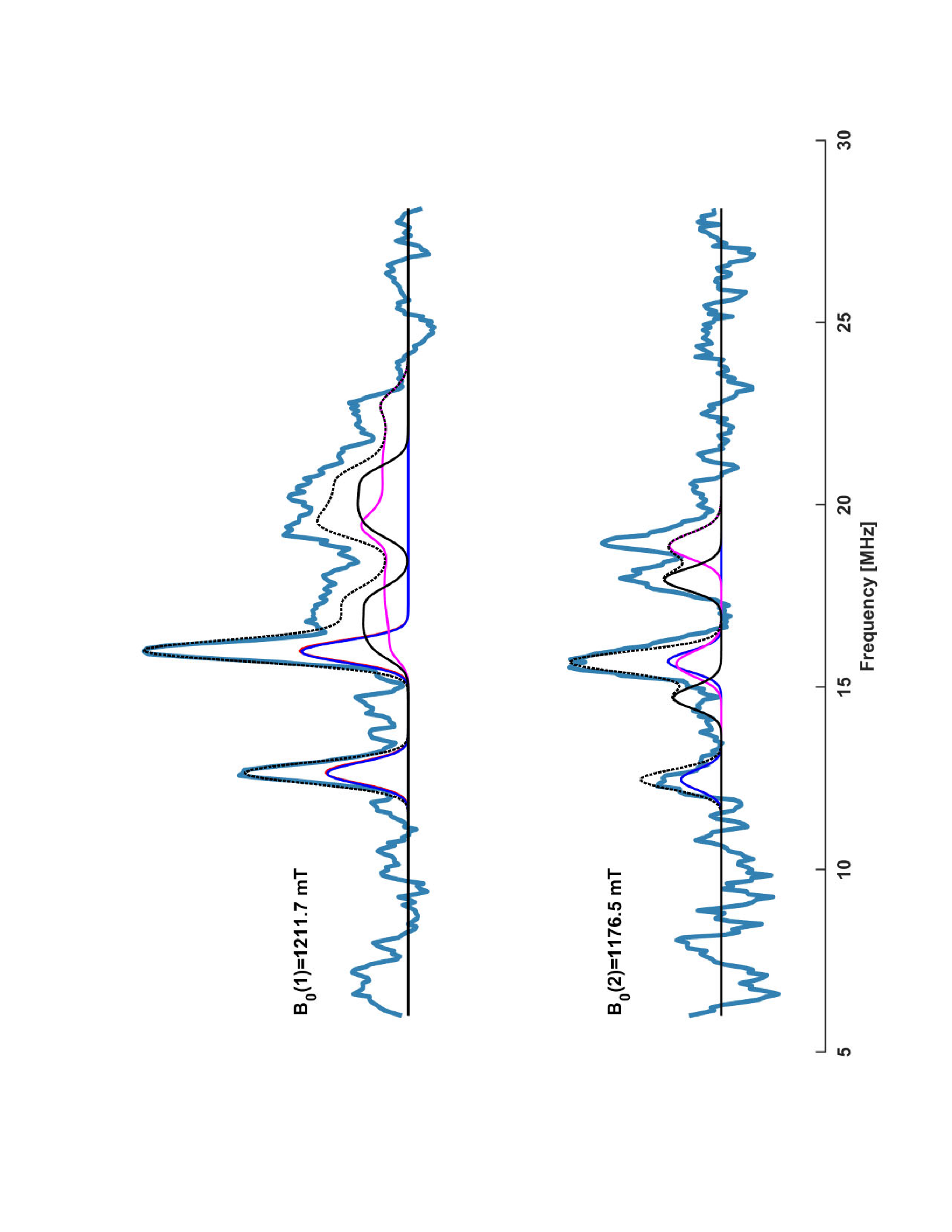

**Figure. S9: ^57^Fe Davies ENDOR of the [Fe_3_-S_4_-Fe-ACP]^3+^ species.** Experimental data (pale blue) shows that the hyperfine contribution of the ^57^Fe in the [Fe_3_-S_4_-Fe-ACP]^3+^ cluster is anisotropic, made evident by its field dependency. Simulations of the nuclear-hyperfine contribution for each of the iron to in the cluster are calculated to Fe_1_ = [28.4, 28.9, 28.1] (red), Fe_2_ = [28.3, 28.9, 28.1] (blue), Fe_3_ = [44, 35, 33.8] (pink), and Fe_4_ = [39.6, 35, 32] (black) the cumulative contribution of each iron simulation is shown as a the black dashed line. Samples were measured at 7 K; MW frequency 33.992 GHz, τ = 184 ns with a [π/2] = 20 ns, and a t[RF] = 26 µs.

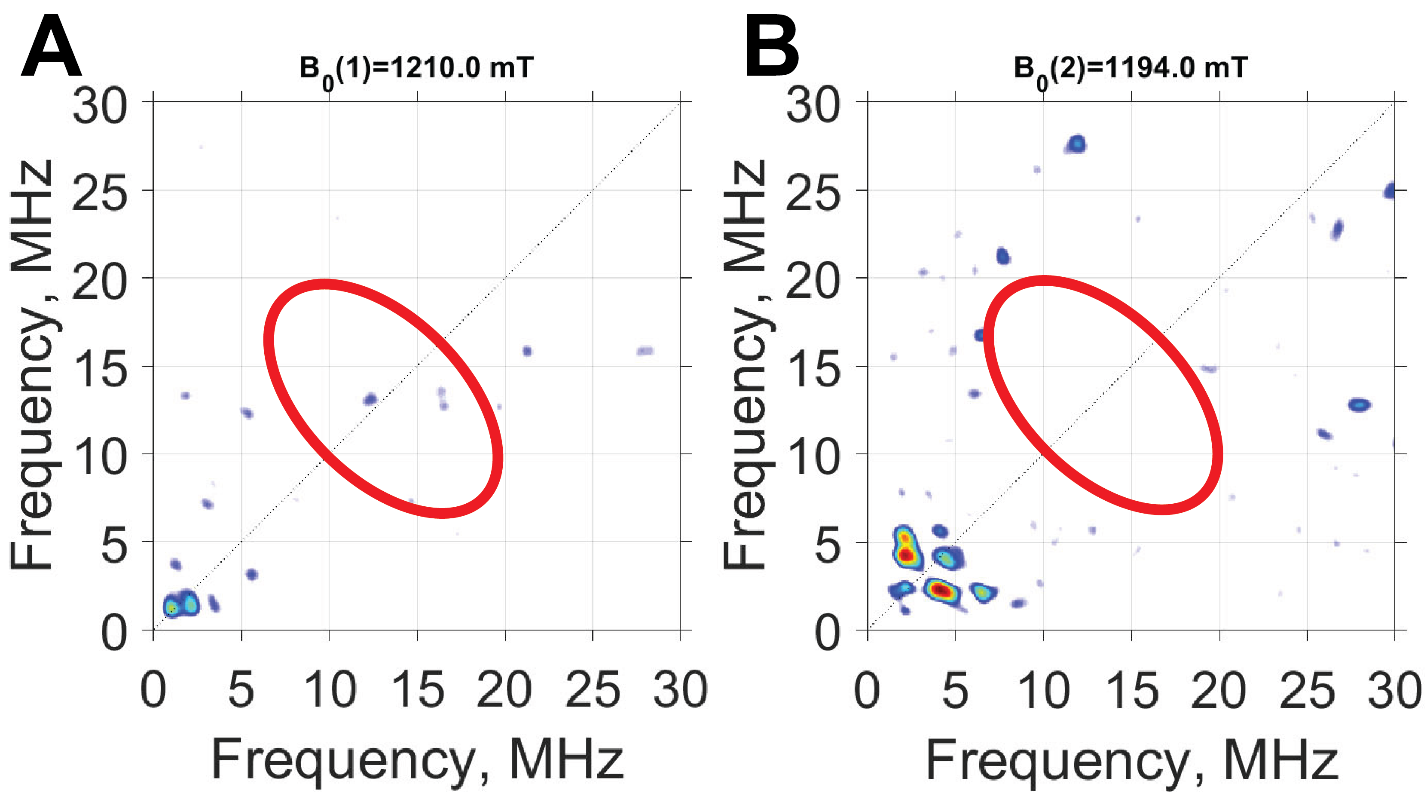

**Figure. S10: HYSCORE of [Fe_3_-S_4_-Fe-ACP]^3+^ with [carboxy-^13^C]-SAM.** A, B) There are no signs of an additional cross-correlation ridge derived from the carboxy-^13^C-labeled carbon of the ACP coupling to the open iron of the cluster. Samples were measured at 8 K at the field positions indicated above the respective plots; MW frequency 34.01 GHz, τ = 192 and 160 ns, and [π/2] = 12 ns.

| **Cluster** | **Structure** | **g-valuses** | | | **g_avg_** | **Carbon Coupling** | **Citation** |
| --- | --- | --- | --- | --- | --- | --- | --- |
| *Pa*ArsL  as-isolated | [Fe_3_S_4_]^1+^ | 1.960 | 1.993 | 2.025 | 1.993 | - | This work |
| *Pa*ArsL reduced | [Fe_4_S_4_]^1+^ | 1.893 | 1.911 | 2.029 | 1.944 | - | This work |
| *Pa*ArsL  SAM bound | [Fe_4_S_4_]^1+^ | - | - | - | - | C1 = 0.22 MHz | This work |
| *Pa*ArsL  SAH bound | [Fe_4_S_4_]^1+^ | 1.872 | 1.905 | 2.011 | 1.929 | - | This work |
| *P. Falciparum* IspH reduced | [Fe_4_S_4_]^1+^ | 1.899 | 1.899 | 2.036 | 1.945 | - | [1] |
| *P. Falciparum* IspH  product bound | [Fe_4_S_4_]^1+^ | 1.933 | 2.010 | 2.084 | 2.009 | 0.2 MHz ≤ a_iso_≤ 3.6 MHz* | [1] |
| *E.Coli* PFL-AE SAM bound | [Fe_4_S_4_]^1+^ | 1.870 | 1.880 | 2.010 | 1.920 | C1 = 0.22 MHz | [2] |
| *Pa*ArsL organometallic | [Fe_3_-S_4_-Fe-ACP]^3+^ | 2.005 | 2.023 | 2.130 | 2.053 | Cγ = 21 MHz | This work |
| *P. Falciparum* IspH Intermediate | [Fe_4_S_4_]^3+^ IPP^-^ | 1.997 | 2.010 | 2.175 | 2.061 | - | [1] |
| *E.Halophila* (HiPIP) | [Fe_4_S_4_]^3+^ | 2.030 | 2.030 | 2.146 | 2.069 | - | [3] |
| *Synechocystis* FTR-NEM | [Fe_4_S_4_]^3+^ | 1.974 | 1.992 | 2.109 | 2.030 | - | [4] |
| *Spinacia*  FTR-NEM | [Fe_4_S_4_]^3+^ | 1.980 | 2.000 | 2.110 | 2.053 | - | [5] |
| Synthetic Alkyl Cluster (methyl) | [Fe_3_-S_4_-Fe-CH_3_]^3+^ | 2.030 | 2.040 | 2.090 | 2.053 | C_methyl_ = 5.5 MHz | [6] |
| Synthetic cluster | [Fe_4_S_4_]^3+^ | 2.030 | 2.050 | 2.100 | 2.060 | - | [7] |
| *T. thermophilus* IspG  Intermediate "X" | [Fe_3_-S_4_-Fe-MECPP]^3+^ | 1.999 | 2.018 | 2.083 | 2.033 | C3 = 3.0 MHz  C2 17.7 MHz | [8] |
| Dph2 Int I | [Fe_3_-S_4_-Fe-ACP]^3+^ | 2.005 | 2.005 | 2.036 | 2.015 | Cγ = 7.8 MHz | [9] |
| Ω  (*E.Coli* PFL-AE) | [Fe_3_-S_4_-Fe-5′dA]^3+^ | 2.004 | 2.004 | 2.035 | 2.014 | C5′ 9.4 MHz  C_methyl_ = 0.5 MHz | [10] |
| *A. aeolicus* ferredoxin | [Fe_2_S_2_]^1+^ | 1.880 | 1.960 | 1.050 | 1.630 | - | [11] |
| *T. thermophilus* ferredoxin | [Fe_2_S_2_]^1+^ | 1.810 | 1.940 | 2.140 | 1.963 | - | [12] |
| * IspH inhibitors were used to get the upper limit | | | | | | | |

**Table S2: FeS Cofactor reference table.**

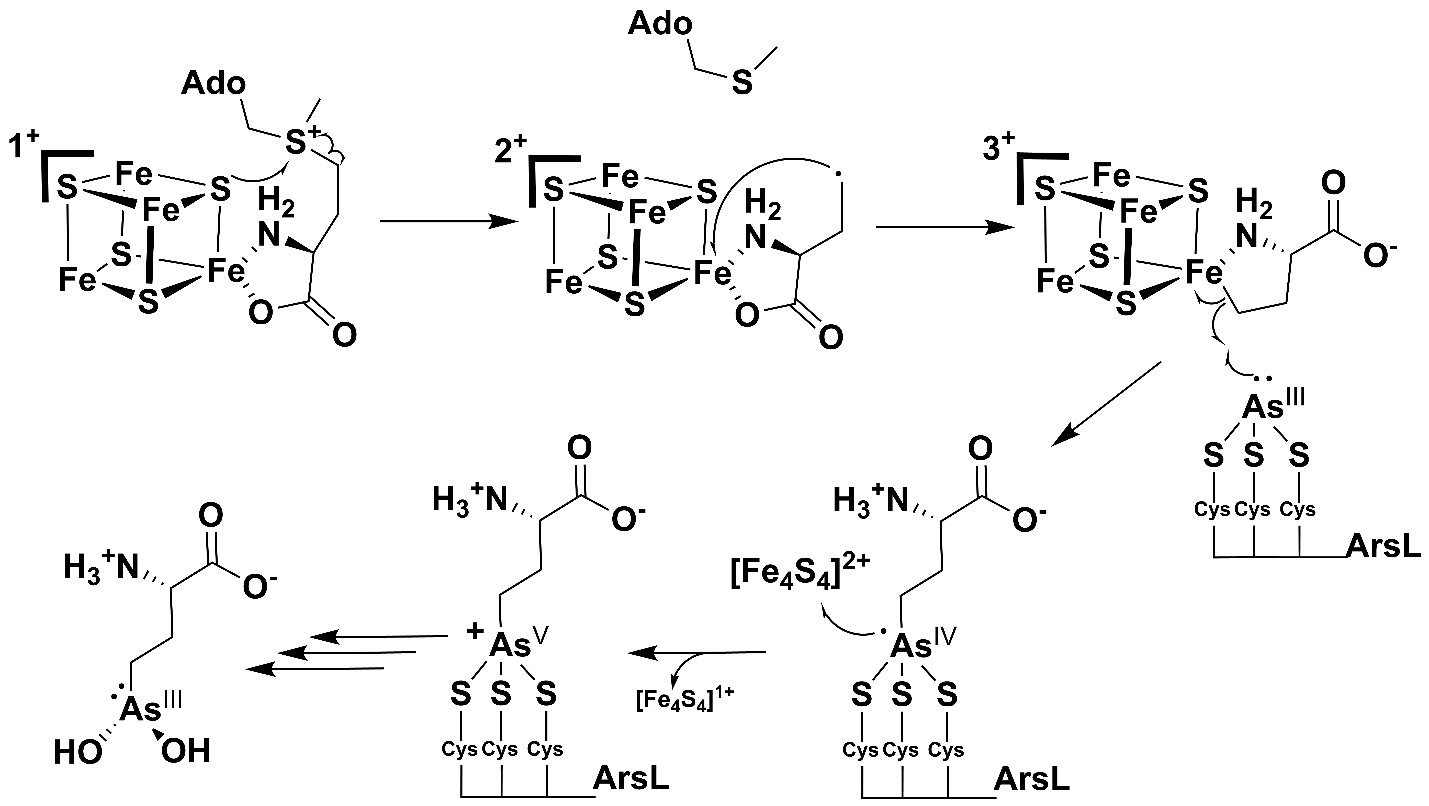

**Scheme. S1: Alternative ArsL reaction mechanism.** In this alternative mechanism, we hypothesize Fe-carbon bond undergoes homolysis, reforming the ACP• to perform a radical attack on the bound arsenous acid. This forms a transient As(IV) AST-OH product-bound intermediate. This As(IV) species then donates a single electron to the [Fe_4_S_4_]^2+^ cluster.
